# Vicennial metagenomic time series unveils evolutionary dynamics of giant viruses in a freshwater ecosystem

**DOI:** 10.1101/2025.06.13.655548

**Authors:** Yumary M. Vasquez, Miguel F. Romero, Robert Bowers, Robin R. Rohwer, Katherine McMahon, Tanja Woyke, Frederik Schulz

## Abstract

Giant viruses represent key ecological players in aquatic ecosystems, yet their evolutionary dynamics in response to environmental change remain poorly understood, particularly in freshwater environments. We leveraged an unprecedented 20-year time series (2000-2019) of 471 co-assembled metagenomes from Lake Mendota (USA) to reconstruct 1,512 high-quality giant virus metagenome-assembled genomes (GVMAGs), providing a unique framework to track viral genome evolution across decades. Viruses in the order *Imitervirales* dominated the Lake Mendota virome, exhibiting consistent presence across all seasons and years. We identified gene duplication (23% of genes) and horizontal gene transfer (29% of genes) as drivers of genomic innovation in giant viruses. Co-occurrence network analysis between viral DNA polymerase B and eukaryotic 18S rRNA sequences revealed significantly increased virus-host associations following the 2009 invasion of a predatory zooplankton. Genome-wide single nucleotide polymorphism analysis demonstrated predominantly purifying selection across viral genes, but revealed a significant increase in positively selected genes post-invasion, including in functions related to host infection (such as protein translation). Comparative evolutionary analyses revealed that giant viruses exhibit genome-wide substitution rates similar to co-occurring bacteria but significantly slower than smaller dsDNA phages, positioning them as evolutionary intermediates with bacterial-like genomic stability but virus-like adaptive capacity. By analyzing a vicennial (20-year) time series, we show that freshwater giant viruses employ sophisticated evolutionary strategies. They broaden their host range without abandoning established partners and maintain stable genomic backbones while rapidly adapting infection-related genes. These dynamics highlight their critical yet previously underappreciated role in freshwater ecosystem dynamics and resilience to environmental change.

## MAIN

Nucleocytoplasmic large DNA viruses (NCLDVs), commonly known as giant viruses and classified within the viral phylum *Nucleocytoviricota*, occupy an evolutionary frontier between viruses and cellular life. With genomes ranging from hundreds of kilobases to over 2.5 megabases, these viruses infect diverse eukaryotes across marine, freshwater, and terrestrial environments, where they significantly impact ecological dynamics (Suttle, 2007; Schulz, Abergel and Woyke, 2022). Notably, many giant virus species maintain their own machinery for DNA repair, transcription, and even translation, challenging traditional boundaries between viral and cellular domains (Raoult *et al*., 2004; Abergel *et al*., 2007; Yutin *et al*., 2013; Schulz *et al*., 2017). Despite their ecological significance and genomic complexity, how these viruses evolve in response to environmental change remains poorly understood, particularly in natural settings where host communities fluctuate over time.

Giant virus genomes exhibit remarkable evolutionary plasticity, with genomic diversity emerging through a balance of gene acquisition and loss, a process termed the “genomic accordion” (Filée, 2015). This expansion and contraction of genomes enables rapid adaptation to environmental pressures while maintaining core viral proteins. Multiple molecular mechanisms drive this genomic innovation: horizontal gene transfer from hosts and other microbes provides “ready-made” functional genes (Filée, Pouget and Chandler, 2008; Moreira and Brochier-Armanet, 2008; Yutin, Wolf and Koonin, 2014), gene duplication creates opportunities for functional divergence (Machado *et al*., 2023), and de novo gene emergence potentially generates entirely novel proteins (Legendre *et al*., 2019). Despite this genomic plasticity, giant viruses maintain an overall pattern of purifying selection that preserves essential functions (Ogata and Claverie, 2007; Legendre *et al*., 2019). However, our understanding of these evolutionary processes remains constrained to laboratory studies which offer mechanistic insights but examine only single host-virus systems over brief timescales (Retel *et al*., 2022; Le Pennec *et al*., 2024), while studies of environmental samples often lack long-term temporal resolution needed to capture adaptation in progress (Meng *et al*., 2023; Fang *et al*., 2025).

Long-term ecological monitoring of natural ecosystems provides the ideal framework for studying how giant viruses evolve in response to environmental change (Roux *et al*., 2017; Laperriere *et al*., 2024; Fang *et al*., 2025). Among such systems, Lake Mendota (Wisconsin, USA) offers exceptional advantages for studying viral adaptation. This eutrophic freshwater lake has been systematically monitored since 1981, establishing one of North America’s most comprehensive ecological time series (Magnuson, Carpenter and Stanley, 2023). Critically, Lake Mendota experienced a major ecological disruption in 2009 with the detection of the invasive predatory zooplankton spiny water flea (*Bythotrephes cederströmii*), though sediment evidence suggests earlier arrival (Walsh, Carpenter and Vander Zanden, 2016), followed by zebra mussel (*Dreissena polymorpha*) invasion in 2015 (Spear *et al*., 2022). The spiny water flea invasion triggered substantial restructuring of the lake’s protistan communities, including documented declines in cryptophyte diversity and relative abundance (Krinos *et al*., 2024). Such protistan community shifts are particularly relevant for giant viruses, as these eukaryotes, along with other eukaryotic groups, represent their primary hosts. The Lake Mendota system thus serves as a natural experiment to investigate how giant viruses respond and adapt to rapid shifts in host community composition, a question with broad implications for understanding virus-host coevolution and ecosystem resilience in the face of environmental change.

Here, we leverage an unprecedented resource for studying viral evolution: a two-decade metagenomic time series from Lake Mendota comprising 471 samples collected between 2000-2019 (Rohwer *et al*., 2025). Using a metagenomic co-assembly approach (Oliver *et al*., 2024), we reconstructed 1,512 non-redundant giant virus genomes, a collection comparable in size to the 1,885 bacterial genomes assembled from the same dataset. This framework enables us to address three fundamental questions about giant virus evolution: (1) How do mechanisms of genomic innovation (*i.e.,* gene duplication, horizontal gene transfer, and de novo gene emergence) operate in natural settings? (2) How does ecosystem disruption by invasive species reshape giant virus-host relationships and the selective pressure acting on giant virus genomes? (3) How do evolutionary rates of giant viruses compare to those of bacteria and smaller viruses in the same ecosystem? Our analyses reveal that the 2009 spiny water flea invasion serves as a natural experiment, allowing us to directly observe genomic adaptations including real-time gene gains and losses, shifts in host association networks, and accelerated selection on infection-related genes. By comparing substitution rates across microbial domains, we demonstrate that giant viruses occupy an evolutionary middle ground, with genome-wide rates matching co-occurring bacteria while adaptive rates more closely resemble other viruses. This work demonstrates that giant viruses respond dynamically to ecological perturbation while maintaining long-term population stability, establishing their importance in freshwater ecosystem resilience.

## RESULTS

### Community structure and temporal distribution of Lake Mendota’s giant viruses

We recovered 1,512 high-quality, non-redundant giant virus metagenomic-assembled genomes (GVMAGs) from Lake Mendota’s 20-year time series (2000-2019), spanning six distinct seasonal periods (Oliver *et al*., 2024; Rohwer *et al*., 2025) (Fig. 1a). This viral genome count approaches the 1,885 bacterial genomes assembled from the same dataset (Oliver *et al*., 2024), highlighting the substantial contribution of giant viruses to freshwater microbiome genome diversity. Taxonomic classification revealed the majority of GVMAGs belonged to the order *Imitervirales* (n = 1,354; 89.6% of all GVMAGs), a pattern consistent with observations from other ecosystems (Ha, Moniruzzaman and Aylward, 2023; Pitot *et al*., 2024; Zhang *et al*., 2024). Within *Imitervirales*, *Mesomimiviridae* constituted the majority (n = 918; 60.7% of all GVMAGs), followed by *Mimiviridae* (n = 197) and *Allomimiviridae* (n = 84). Other orders, *Pimascovirales* (n = 115), *Asfuvirales* (n = 26), *Pandoravirales* (n = 8), and *Algavirales* (n = 9), were present but less abundant. Only PM_01 (order *Pimascovirales*, n = 83) exceeded 50 genomes among non-*Imitervirales* families (Supplementary Table S1). The predominance of *Mesomimiviridae* in Lake Mendota mirrors their dominance in marine systems (Aylward *et al*., 2021), suggesting conserved ecological advantages for this family across aquatic environments.

**Figure 1:**
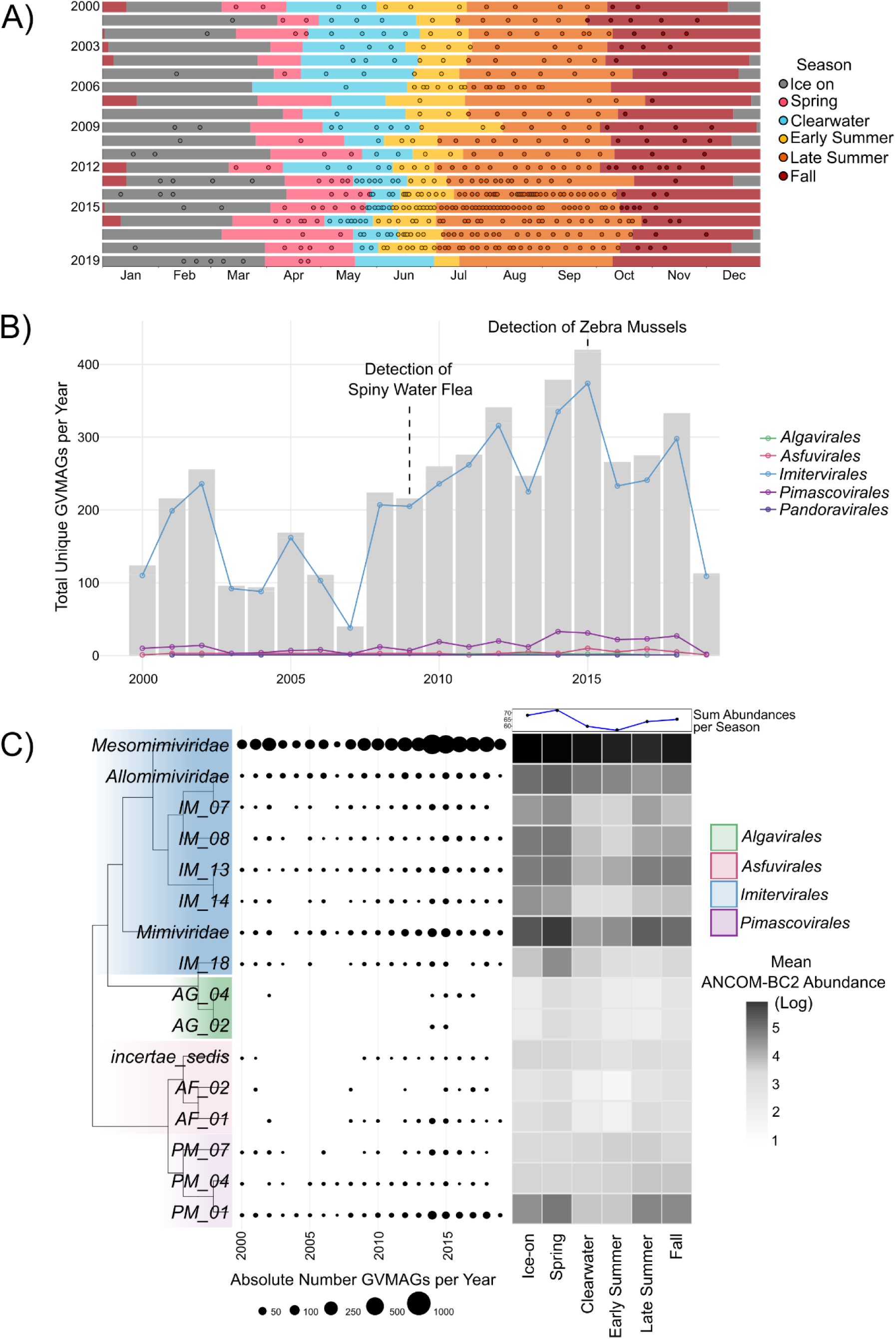
Giant virus population dynamics in Lake Mendota. A) Distribution of metagenome samples from Lake Mendota across 20 years and 6 seasons. Open black circles denote the sampling time points for each metagenome sample. B) Number of unique giant viruses with at least 1x coverage and 20% of the genome mapped *(i.e.*, breadth) mapped per year (barplot) with the breakdown of composition by family (line plot). C) Giant virus presence in each year and season broken down to family level. Years include the total number of giant virus MAGs (not unique) per year with at least a mean coverage of 1 and at least 20% breadth mapped per year. Season presence includes the log transformation of the mean abundance calculated using ANCOM-BC2 which takes into account the number of samples per year.

Tracking GVMAGs across the 20-year time series revealed distinctive temporal persistence patterns (Fig. 1b,c). *Imitervirales* maintained consistent presence throughout all 20 years, establishing this order as the stable foundation of Lake Mendota’s giant virus community. In contrast, other orders displayed intermittent detection: *Algavirales* appeared briefly in 2002, became undetectable for over a decade, then reemerged during 2014-2017, while *Asfuvirales* was detected during 2000-2002 and later during 2008-2019. The single *Algavirales* GVMAG detected in 2002 was not observed again, suggesting three possible scenarios: adaptation of existing viral strains to changing conditions, emergence of new viral strains, or methodological limitations in years with lower sampling intensity (*i.e.,* less deep sequencing). We also identified clear seasonal dynamics (Fig. 1c), with bias-corrected abundance analysis (ANCOM-BC2) revealing that giant virus abundance peaks during Spring, reaches its minimum in Early Summer, and gradually increases through Fall and Ice-on before reaching another Spring maximum. This seasonal pattern closely tracks eukaryotic host abundance cycles, with microeukaryotic taxa reaching maximum abundance during Spring, Ice-on, and Fall periods (Krinos *et al*., 2024). The Early Summer minimum follows the lake’s "Clearwater" phase, where phytoplankton abundance and biovolume are low (Carey *et al*., 2016). The tight coupling between giant virus and eukaryotic seasonal patterns supports the expected ecological relationship where viral abundance follows host availability.

We validated our co-assembly approach through comprehensive methodological comparisons with traditional single-sample assemblies. Multiple statistical tests confirmed that both methods captured nearly identical seasonal community dynamics. ANCOM-BC2 analysis demonstrated that only three viral families (AG_02, PM_07, PV_02) showed significant interaction effects between seasons and assembly method; for all other families, seasonal patterns remained consistent regardless of assembly approach (Supplementary Figures S1 & S2). Direct ordination comparison through Procrustes analysis revealed exceptional congruence between methods (r = 0.96, p = 0.003) (Supplementary Figure S3). This finding was further supported by strong Mantel correlations between seasonal dissimilarity matrices (r = 0.89, p = 0.0014). The robust correlation persisted even after compositional data transformation (center log-ratio; r = 0.75) and nearly identical magnitudes of seasonal transitions (r = 0.87), though with reduced statistical significance (p = 0.068; p = 0.054, respectively). These multiple lines of evidence confirm that our co-assembly approach reliably captures seasonal giant virus dynamics, differing from single-assembly results only by a slight consistent offset and modestly increased variability during summer months.

### High ortholog conservation suggests common selective pressures shape giant virus genomes

Lake Mendota giant virus genomes exhibit remarkably high genetic conservation despite their taxonomic diversity. Our ortholog analysis revealed that approximately 86% of all genes are shared between multiple virus species, while only 14% are species-specific or unassigned to orthogroups. At least half of each giant virus genome falls within an orthogroup (Supplementary Figure S4). Only a single genome, of uncertain taxonomy but potentially belonging to *Pandoravirales*, showed lower conservation (41.7% genes in orthogroups). The degree of gene sharing varies significantly across viral families, *Imitervirales* members display the highest proportion of conserved genes, while *Asfuvirales* show the lowest (Supplementary Figure S5). While sampling bias partially explains this pattern, the abundant *Imitervirales* genomes (n=1,354) naturally share more orthogroups than the rarer *Asfuvirales* (n=26), the overall level of gene sharing is comparably consistent (83% reported for 207 NCLDV genomes; Sun and Ku, 2021). Previous studies comparing giant viruses across diverse environments typically report lower inter-order gene conservation (Schulz *et al*., 2020). This unusually high functional overlap likely reflects shared selective pressures within Lake Mendota’s relatively homogeneous environment, where similar eukaryotic host communities and physicochemical conditions impose convergent functional constraints (Sun and Ku, 2021). Unlike viruses adapting to radically different habitats, Lake Mendota’s giant viruses appear to have converged on similar genetic solutions to common ecological challenges, creating a functionally cohesive viral community despite their taxonomic diversity.

We reclustered the predicted proteins from all Lake Mendota giant viruses together with those of 630 marine giant virus genomes from the Tara Oceans sequencing effort, yielding 31,222 orthogroups and 98,952 singletons. More than half of the orthogroups (n = 17,348; 55.5%) and singletons (n = 72,032; 73%) are populated exclusively by proteins found in Lake Mendota genomes, whereas only 4,967 orthogroups (15.9%) and 26,920 singletons (37%) are unique to the Tara Oceans genomes. The remaining 8,907 orthogroups (28.5%) contain sequences from both environments. Thus, although almost one-third of orthogroups encode functions common to freshwater and marine viruses, the majority appear ecosystem-specific, reinforcing the idea that the ecological challenges present in Lake Mendota have led to a shared genetic toolkit.

### Duplication, acquisition, and real-time genome restructuring in giant viruses

Gene gain, particularly through gene duplication, is a major force driving the expansion of giant virus genomes (Machado et al., 2023). Our analysis revealed that species-specific gene duplications constitute a substantial portion of giant virus genetic material, accounting for approximately 23% of all genes across our dataset. The prevalence of duplicated genes varies systematically across taxonomic levels, suggesting clade-specific evolutionary strategies. At the order level, *Pimascovirales* genomes showed the highest duplication rates (25% of genes), while *Pandoravirales* exhibited the lowest (19%). These patterns become more pronounced at the family level, with certain families (*e.g.*, *Algavirales* families AG_04 and AG_02) maintaining notably lower duplication rates compared to other viral lineages (Fig. 2a). The variation becomes even more pronounced at the individual genome level, where the proportion of duplicated genes ranged dramatically from 6% to 60%. This ten-fold difference suggests fundamentally different evolutionary regimes operating across giant virus lineages, potentially reflecting diverse selective pressures, host adaptation strategies, or intrinsic differences in genome replication fidelity.

**Figure 2:**
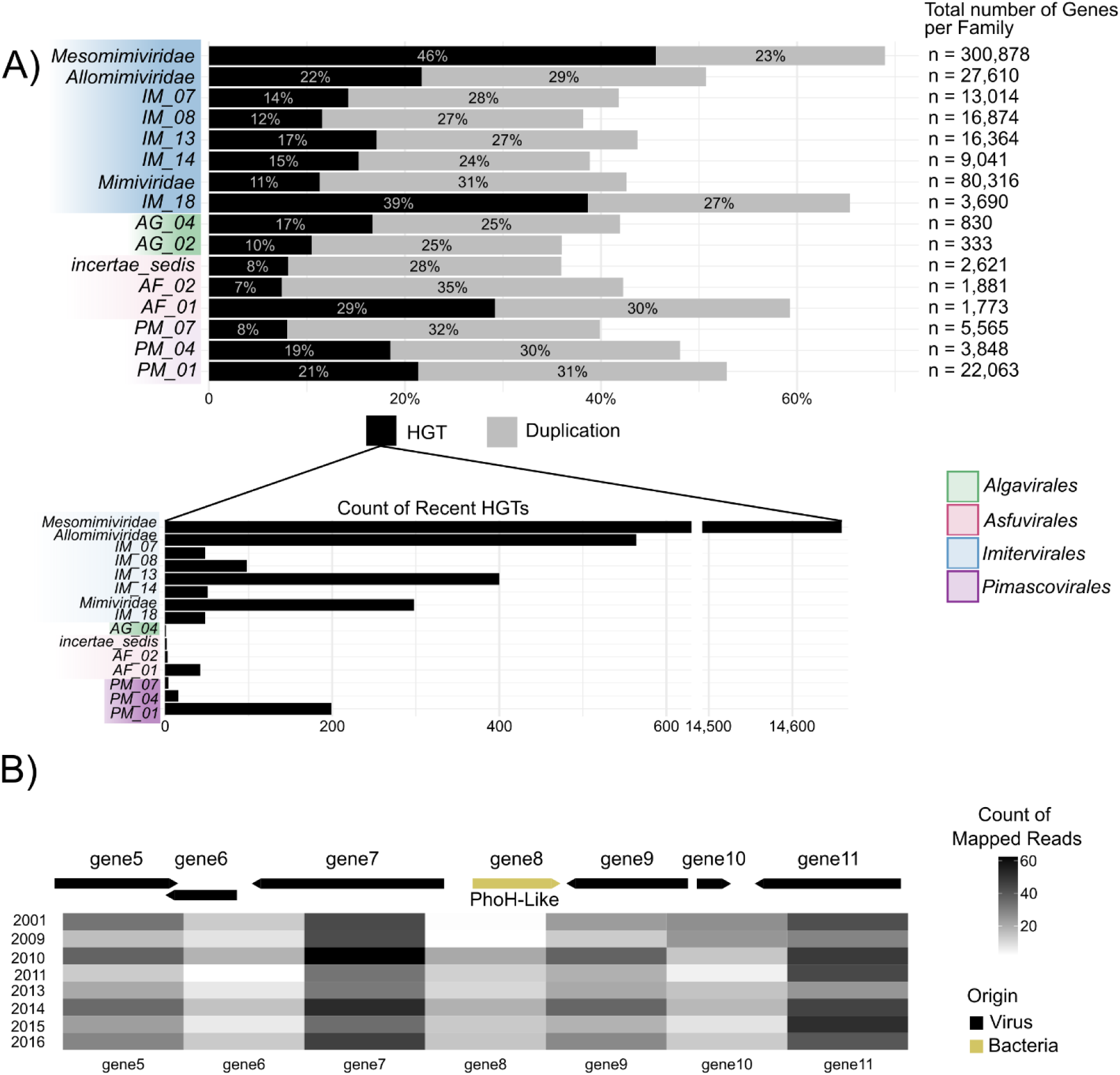
Horizontal gene transfer (HGT) and duplications and in giant viruses. A) Percentage of giant virus genes per viral family that are predicted to be the result of horizontal gene transfer or gene duplication. HGTs are further broken down to those that have a high percent identity to a non-viral homolog. B) Gene gain in a *Mesomimiviridae* genome following spiny water flea invasion in 2009 (See Supplementary Figure S14 for information on years 2002 - 2008). Sparse and inconsistent mapping was observed in years prior to 2009 (2002–2008), with no reliable counts obtained; these data are not shown. BLASTp results indicate a homolog to the PhoH gene in bacteria. Total mapped reads per gene are indicated by the gradient colors.

Functional analysis of duplicated genes reveals expansion of certain viral capabilities. While most duplicated genes lack matches to characterized functions in the KEGG database, the annotated fraction shows expansion in four primary categories: post-translational modification, protein turnover, and chaperone functions (COG group O; n = 8,147 genes); DNA replication and repair (L; n = 7,913); cell wall, membrane, and envelope biogenesis (M; n = 6,155); and transcription (K; n = 4,150). This pattern indicates that duplication events preferentially amplify core viral functions already prevalent across *Nucleocytoviricota* (Yutin and Koonin, 2012; Karki, Moniruzzaman and Aylward, 2021), potentially enhancing infection efficiency. For example, we identified duplications in genes involved in ubiquitination (K08770), which are thought to be involved in host protein degradation or in host signaling (Boyer *et al*., 2009; Moniruzzaman, Gann and Wilhelm, 2018). Whether these duplicated genes are truly functional in all Lake Mendota GVMAGs, or simply represent genomic redundancy, remains unclear and will require future transcriptomic or proteomic studies. Nevertheless, our findings strongly suggest that gene duplication serves as a key mechanism driving genome expansion in giant viruses and may significantly contribute to their ecological success and evolutionary diversification across aquatic environments.

Horizontal gene transfer (HGT) represents an equally significant driver of giant virus genome innovation, accounting for 29% of all genes in our dataset, comparable to gene duplication (23%) but with distinct taxonomic and functional patterns. Notably, 8% of genes showed evidence of both duplication and HGT. Using stringent identity thresholds against custom Lake Mendota bacterial assemblies and the NR database (≥50% identity for historical transfers; ≥80% for recent transfers), we identified taxonomic differences in HGT frequency. *Pandoravirales* genomes contained remarkably few horizontally acquired genes (3%), while *Imitervirales* showed ten-fold higher HGT rates (32%) (Fig. 2a). This pattern mirrors observations in viral-viral gene transfer studies (Wu *et al*., 2024), suggesting that lineage-specific factors may shape the likelihood for horizontal gene acquisition regardless of the gene’s origin. Functionally, HGTs were non-randomly distributed, with significant enrichment in replication and repair genes (L; n = 14,707) and transcription (K; n = 9,969), critical functions that aid viral reproduction capacity. These findings align with previous studies documenting non-viral origins of many giant virus DNA repair genes (Ogata *et al*., 2011; Redrejo-Rodríguez and Salas, 2014; Blanc-Mathieu and Ogata, 2016). Particularly interesting are the families IM_13 and IM_14, where recent HGTs (≥80% identity) constitute 18% and 17% of all horizontally acquired genes respectively (Fig. 2a). This recent acquisition of genes into specific viral families suggests that horizontal gene transfer represents an ongoing mechanism shaping giant virus evolution.

Our 20-year time series provides a rare opportunity to directly observe real-time gene gain events in giant viruses. Using stringent criteria to minimize false positives, we identified 17 putative gene gains across 10 genomes, predominantly in *Imitervirales* (seven *Mesomimiviridae*, one IM_13, one IM_07) with a single event in *Pimascovirales* (PM_01). Our methodology required both flanking gene presence to exclude mapping artifacts and temporal separation between virus detection and gene acquisition to ensure true evolutionary events rather than assembly or detection artifacts. Gene duplication overwhelmingly dominated the gain landscape, accounting for 16 of 17 events, reinforcing the central role of duplication in giant virus genome evolution. Eight gained genes had homologs in the NCBI non-redundant database or Lake Mendota bacterial MAGs, while only three could be functionally characterized: a nematode hypothetical protein, a DNA-directed RNA polymerase, and a phosphate starvation-inducible protein (phoH) (Fig. 3b). The phoH acquisition is particularly significant as this gene may help giant viruses cope with nutrient-limited conditions by supporting energy homeostasis during phosphate starvation (Moniruzzaman *et al*., 2020). Intriguingly, all three functions existed in other Lake Mendota giant viruses but were previously absent in the recipient genomes, indicating selective acquisition of functions with proven utility in the lake environment. The timing of gene gains showed potential correlation with ecological events, with six gains occurring in 2009-2010, coinciding with the spiny water flea invasion and subsequent protistan community restructuring. We identified one gene (921 bp, 40.1% GC) lacking any homologs or duplication signature, potentially representing de novo gene emergence. This gene shows high structural confidence (pLDDT = 74.06), but minimal sequence identity (28%) to an uncharacterized protein from a marine bacterium (Jumper *et al*., 2021). The presence of this putative de novo gene highlights the potential for giant viruses to generate novel genetic material, though its functionality and expression remain to be determined.

**Figure 3:**
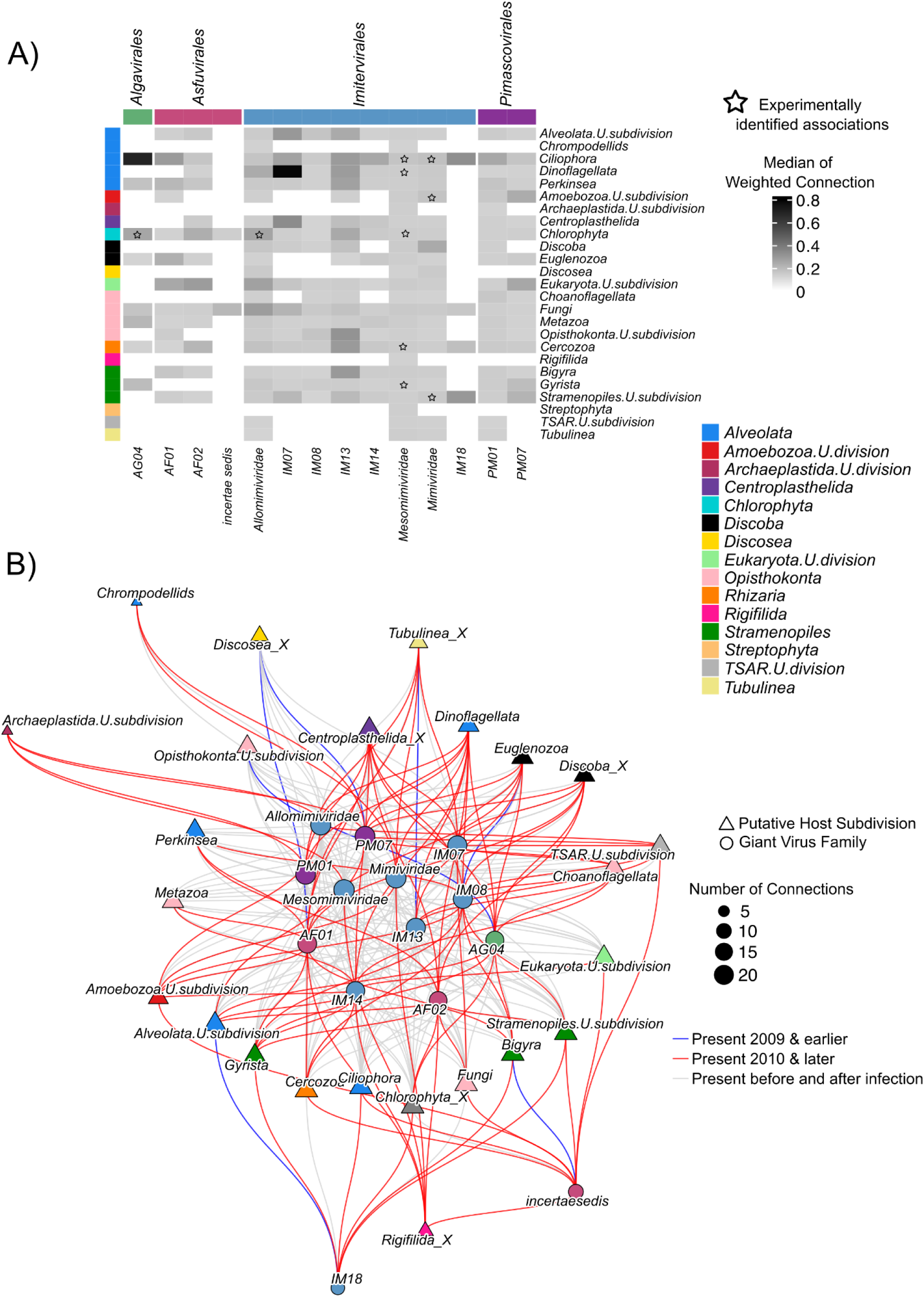
Putative eukaryotic hosts of giant viruses. A) Weighted connection of *Nucleocytoviricota*-eukaryote pairs that takes into consideration the number of consecutive years where the association occurs. Stars indicate experimentally confirmed eukaryote-giant virus associations from published literature. B) Network associations before and after the invasion of the spiny water flea in Lake Mendota. Associations present before and after infections are colored in grey, associations present in and before 2009 colored in blue, and associations formed in 2010 and later are colored in red. Eukaryotic host subdivision (triangle) and giant virus family (circle) are colored by their group color as seen in A, as sized by the number of edges (connections).

Parallel to gene gains, we documented 13 gene loss events across 12 viral genomes, suggesting a dynamic balance between genome expansion and contraction. The majority of losses occurred in *Imitervirales* (10 in *Mesomimiviridae*, one in *Allomimiviridae*), with a single event in an unclassified genome within class Megaviricetes. Unlike gene gains, which showed temporal gene events in 2009-2010, we observed no gene losses during these years. Eight lost genes had identifiable homologs in the NR database or Lake Mendota bacterial MAGs, while only three could be functionally characterized: ribonuclease H, exocyst complex component sec10, and a gene annotated as "determination of stomach left/right asymmetry." The functional implications of these losses appear minimal; while ribonuclease H is widely distributed across giant virus genomes and likely functionally important, the other two genes have restricted distribution even within our dataset, suggesting their loss may have limited impact on overall viral function. Most lost genes (n = 11) belonged to established orthogroups, indicating they had homologs in other giant viruses, including one paralog that represented a previously duplicated gene. The remaining two lost genes were ORFans unique to single genomes, with one showing homology to Lake Mendota bacterial MAGs, potentially representing a horizontally acquired gene that proved non-adaptive. The final lost gene (297 bp, 50% GC) lacked matches in any reference database or among other Lake Mendota viruses, but AlphaFold analysis revealed moderate sequence similarity (83%) to an uncharacterized protein from a Lake Croche (Canada) viral metagenome, albeit with low structural confidence (pLDDT = 54.5). This particular gene loss suggests that while the gene may share sequence similarity with another freshwater viral protein, its low structural confidence and absence from other freshwater genomes indicate it was likely non-essential and subject to rapid loss. The observed pattern of gene losses, particularly of genes with redundant or poorly characterized functions, supports the concept that genome gigantism in these viruses is maintained through a dynamic balance of gene acquisition and turnover, allowing essential functions to be preserved even as individual genes are lost (Alempic *et al*., 2024). Through this process, giant viruses can rapidly adapt to changing conditions without compromising essential viral processes.

### Virus-host relationships restructure but persist following predator invasion

The 2009 invasion of the spiny water flea (*Bythotrephes cederströmii*) in Lake Mendota created a natural experiment to investigate how giant viruses respond to ecosystem disruption through predicted shifts in host associations. To study these dynamics, we conducted a co-occurrence analysis between 18,375 viral DNA polymerase B (PolB) sequences and eukaryotic 18S rRNA genes across our 20-year metagenomic time series. We selected PolB as our marker gene because it is highly conserved across all *Nucleocytoviricota* while providing sufficient variation for reliable taxonomic classification. While this approach builds on established virus-host prediction methodologies, there are inherent limitations: co-occurrence networks suggest potential infection relationships, but do not prove direct evidence, which would require experimental validation. Nevertheless, this network analysis allows us to predict how viral host range might respond to major ecological disturbances.

Our final co-occurrence network comprised 7,423 *Nucleocytoviricota* PolB sequences and 4,561 eukaryotic 18S rRNA gene sequences, with positive associations dominating the network (96.5%) compared to negative associations (3.5%) (Supplementary Table S2). Network connectivity varied substantially across viral families: *Mesomimiviridae* showed exceptionally high connectivity (consistent with their genomic dominance in the lake), followed by *Allomimiviridae*, *Mimiviridae*, and PM_01 (Supplementary Table S2). Host associations clustered within three major eukaryotic divisions: *Alveolata*, *Opisthokonta*, and *Stramenopiles*, with the TSAR supergroup (*Stramenopiles*, *Alveolata*, and *Rhizaria*) accounting for over half of all detected relationships (Supplementary Table S3). Other major associations included *Obazoa* lineages (particularly *Fungi*, *Metazoa*, and *Choanoflagellata*), *Archaeplastida* (*Chlorophyta*), and various unclassified eukaryotes. This taxonomic distribution broadly aligns with known giant virus host ranges while revealing potential novel associations requiring experimental confirmation (Sun *et al*., 2020).

Each viral family exhibited distinct host association patterns. *Mesomimiviridae* showed diverse host associations dominated by relationships with *Alveolata* (particularly *Ciliophora*; n = 1,961 edges) and *Opisthokonta* (particularly Fungi; n = 1,353 edges). The strong fungal associations observed across all viral families supports recent metagenomic evidence for fungal infection by giant viruses (Bhattacharjee *et al*., 2023; Myers *et al*., 2024), yet contrasts with traditional isolation-based studies that have yet to recover fungi as giant virus hosts. Weighted connection score analysis revealed that *Mesomimiviridae* lacks strong preference for any single eukaryotic group (Fig. 3a), displaying instead a generalist strategy with connections across diverse protistan lineages. This broad host range aligns with previous reports characterizing *Mesomimiviridae* as ecological generalists capable of infecting taxonomically diverse hosts (Sun *et al*., 2020; Meng *et al*., 2021; Sun and Ku, 2021). Phylogeny-aware enrichment analysis further confirmed statistically significant associations between *Mesomimiviridae* and multiple eukaryotic lineages including *Ciliophora*, *Euglenozoa*, *Oomycota*, *Apicomplexa*, *Ascomycota*, and *Chlorophyta* (Supplementary Figure S6), validating our network-based approach.

Temporal dynamics analysis revealed immediate coupling between host availability and viral abundance, with cross-correlation and lagged regression analyses showing strongest relationships at lag 0 (same-month correlation). We found only minor correlations at positive monthly lags, suggesting limited delayed viral responses to host abundance changes (Supplementary Figure S7). This tight temporal coupling indicates rapid viral replication following host availability, consistent with the short infection cycles typical of giant viruses. To isolate the specific impact of the spiny water flea invasion on viral community composition, we performed a distance-based redundancy analysis (db-RDA) while controlling for seven key environmental variables (water temperature, dissolved oxygen, oxygen saturation percentage, dissolved inorganic carbon, total inorganic carbon, dissolved organic carbon, and total organic carbon). This multivariate approach confirmed that invasion status significantly influenced viral community structure independent of background environmental variation (Supplementary Table S4). While invasion status explained a relatively small proportion of overall community variation (0.7%), this effect was statistically significant and ecologically meaningful given the complex suite of factors influencing viral communities. This finding provides statistical support for the hypothesis that ecosystem restructuring following invasion, rather than coincidental environmental changes, drove shifts in virus-host association patterns

To investigate how the spiny water flea invasion reshaped virus-host dynamics, we compared association networks between pre-invasion (2000-2009) and post-invasion (2010-2019) periods, focusing on positive associations that represent potential infection relationships (Fig. 3b; Supplementary Figures S8 & S9). We first examined whether the invasion caused an abrupt structural shift in host-virus networks by analyzing annual network metrics (edges, density, modularity) and implementing formal breakpoint analyses. Results revealed no statistically significant sudden shifts in network structure, indicating that ecological impacts developed progressively rather than as a discrete event (Supplementary Figures S8 & S9). Segmented regression identified a statistical breakpoint near 2009, but trends before and after this point lacked statistical significance, further supporting gradual rather than abrupt ecological reorganization (Supplementary Figure S10). Despite this gradual transition, we maintained 2009 as our analytical boundary for three reasons: (1) it corresponds to the first documented detection of the spiny water flea in Lake Mendota (Magnuson, Carpenter and Stanley, 2023), (2) it aligns with invasion ecology theory, which predicts that ecosystem effects emerge progressively as invader populations establish and native communities respond (Simberloff and Gibbons, 2004), and (3) it enables comparison between two ecologically distinct states: before and after invasion establishment. This approach acknowledges the continuous nature of ecological change while facilitating statistical comparison between fundamentally different ecosystem configurations.

Comparing pre-invasion (2000-2009) and post-invasion (2010-2019) networks revealed a restructuring of virus-host association patterns, with a marked increase in total network associations following invasion establishment (Supplementary Table S5). This finding raised an important methodological question: could the observed increase simply reflect improved detection power from deeper sequencing in later years? To address this concern, we implemented a mixed-effects logistic model that explicitly included yearly sequencing effort as a covariate. This rigorous statistical approach confirmed that the probability of virus-host associations remained significantly higher in the post-invasion period even after controlling for sequencing depth (p = 0.046). We further validated this finding through multiple additional analyses, including degree-preserving null models, network rarefaction, removal of highly-connected nodes, and permutation tests (see Supplementary Methods). As a final confirmation, we implemented a bootstrapping approach that identified 72 high-confidence associations in the pre-invasion network versus 99 in the post-invasion network, a 38% increase (Supplementary Figure S11). This consistent signal across multiple analytical approaches provides strong evidence that the observed network expansion represents a genuine ecological response rather than a methodological artifact.

Of the 99 high-confidence associations in the post-invasion network, 46 represented entirely new relationships that emerged following ecosystem disruption (present in ≥95% of bootstrap iterations). For example, the eukaryotic group *Dinoflagellata*, whose abundance increased following the invasion (Krinos *et al*., 2024), formed new associations with multiple viral families (PM_01, IM_13, and *Mimiviridae*). This pattern suggests opportunistic viral exploitation of newly abundant potential hosts. Many moderate-confidence associations (bootstrap confidence 0.50-0.95) from the pre-invasion period strengthened post-invasion, suggesting that previously tentative virus-host relationships became more stable following community restructuring. While 10 high-confidence associations from the pre-invasion period disappeared (Supplementary Figure S12), most retained moderate confidence levels (0.557-0.937) rather than disappearing completely, indicating relationship shifts rather than complete host abandonment. Importantly, 62 high-confidence associations (86% of pre-invasion relationships) persisted across both periods, demonstrating stability in core virus-host relationships despite ecosystem perturbation. Collectively, these findings reveal that giant viruses appear to maintain established infection networks while opportunistically expanding into newly available host niches created by community restructuring. This adaptive approach would enhance viral resilience to environmental change by balancing stability with opportunity, potentially explaining how giant viruses maintain long-term persistence while responding to ecosystem fluctuations.

### Genomic signatures of selection shift following ecosystem disruption

To investigate real-time molecular adaptation in giant viruses, we analyzed fine-scale genetic variation across our 20-year time series. From our 1,512 GVMAGs, we identified 553 (∼37%) genomes that met stringent criteria for reliable single nucleotide polymorphism (SNP) analysis: ≥93% read identity and ≥10× average sequencing depth across multiple timepoints. While relaxing these thresholds (to 90% identity or 5× depth) would have included more genomes, this increased the risk of mismapped reads and false-positive SNPs under less stringent criteria (Supplementary Figure S13). This conservative approach prioritized analytical confidence over maximizing genome inclusion, ensuring that our downstream evolutionary analyses captured genuine biological signals rather than methodological artifacts. The resulting high-confidence genome set spans all major viral families, providing a robust foundation for investigating selective pressures across taxonomically diverse giant viruses.

Across the 553 genomes, we observed a predominant signature of purifying (negative) selection (ratio of nonsynonymous to synonymous substitutions (dN/dS) < 1). This genome-wide pattern confirms previous findings that giant viruses primarily experience selection that preserves protein function (Ogata and Claverie, 2007). Despite this predominant pattern of purifying selection, we detected positive selection (dN/dS > 1) in 328 genomes. These adaptive signals were unevenly distributed taxonomically: *Allomimiviridae* showed the highest proportion of genomes with positive selection (32% of all *Allomimiviridae* genomes), followed by IM_12 (29%), *Mesomimiviridae* (22%), *Mimiviridae* (15%), and PM_01 (19%), while remaining families had fewer than 10 genomes with positively selected genes (Supplementary Table S7). The intensity of positive selection varied between individual genomes, with the proportion of genes under positive selection ranging from just 0.08% to 27%. This variation suggests fundamentally different selective pressures across viral lineages.

To systematically identify drivers of genomic microdiversity, we implemented a negative-binomial mixed model incorporating viral taxonomy, temporal factors, and ecological variables. The model included random effects for viral families and individual genomes, with variance components (family = 0.26; GVMAG = 1.55) revealing that genetic diversity is structured hierarchically: while family-level factors influence SNP accumulation rates, genome-specific characteristics account for approximately six times more variance. This suggests that while broad taxonomic affiliation shapes evolutionary trajectories, individual genome characteristics (*i.e.,* host range, infection dynamics, and/or replication mechanisms) play the dominant role in determining evolutionary rates. Time-series analysis using 2002 as a baseline (selected for adequate sampling and pre-invasion stability) revealed minimal year-to-year variation in SNP rates, with only 2012 showing marginally lower rates (p = 0.048) and 2019 showing significantly higher rates (p = 0.0007, though this finding requires cautious interpretation due to limited sampling). Seasonal patterns emerged more strongly, with SNP counts peaking during Ice-on (p < 0.001) and Spring (p = 0.002) periods relative to Early Summer, while declining significantly in Late Summer (p = 0.001). These seasonal fluctuations suggest that viral evolutionary rates may track seasonal dynamics in host availability. The balance between selection types remained stable throughout the time series, with negatively/neutrally selected genes consistently outnumbering positively selected genes (p < 0.001) and no significant temporal shifts in this balance (interaction terms: p > 0.05).

While year-to-year SNP patterns showed limited variation, we hypothesized that the spiny water flea invasion might have triggered broader shifts in evolutionary dynamics. Comparing pre-invasion (2000-2009) and post-invasion (2010-2019) periods revealed a pattern: after normalizing for sequencing effort, SNP rates were approximately 33% higher in the post-invasion period (p = 0.047). This finding raised the important methodological question similarly raised above: could this increase simply reflect improved detection from deeper sequencing in later years? To address this concern, we implemented multiple analytical safeguards. First, we rarefied the data to ensure identical sequencing depth between periods, confirming that the post-invasion increase remained robust (mean estimate = -0.44; 95% empirical interval = -0.54 to -0.33). Second, we conducted change-point analysis, which supported 2009 as a biologically meaningful breakpoint; an AIC sweep showed minima between 2009-2011, with all models in this range statistically equivalent (ΔAIC < 3). Third, we built multivariate models controlling for key environmental parameters (temperature, oxygen, and carbon chemistry variables), finding that invasion status remained a significant predictor of SNP rates independent of these factors (p < 0.001). Within these models, dissolved inorganic carbon and total organic carbon showed positive associations with SNP counts (p = 0.029 and p = 0.012, respectively), while total inorganic carbon showed negative associations (p = 0.037). These relationships suggest a plausible mechanistic link through carbon availability: viral SNPs arise during replication, and elevated dissolved inorganic carbon and total organic carbon stimulate primary production (Jansson, Karlsson and Jonsson, 2012), increasing protist biomass and turnover. A larger, faster-growing host pool supports more frequent or larger viral outbreaks, creating additional replication cycles and therefore a larger pool of detectable mutations (Knowles *et al*., 2020). When examining viral families with SNP data from both periods, we observed consistent increases in the proportion of genes under selection post-invasion (Fig. 4), with family IM_18 representing the only exception to this pattern. This genome-wide acceleration of selection suggests a broad evolutionary response to ecosystem reorganization rather than simply enhanced mutation detection.

**Figure 4:**
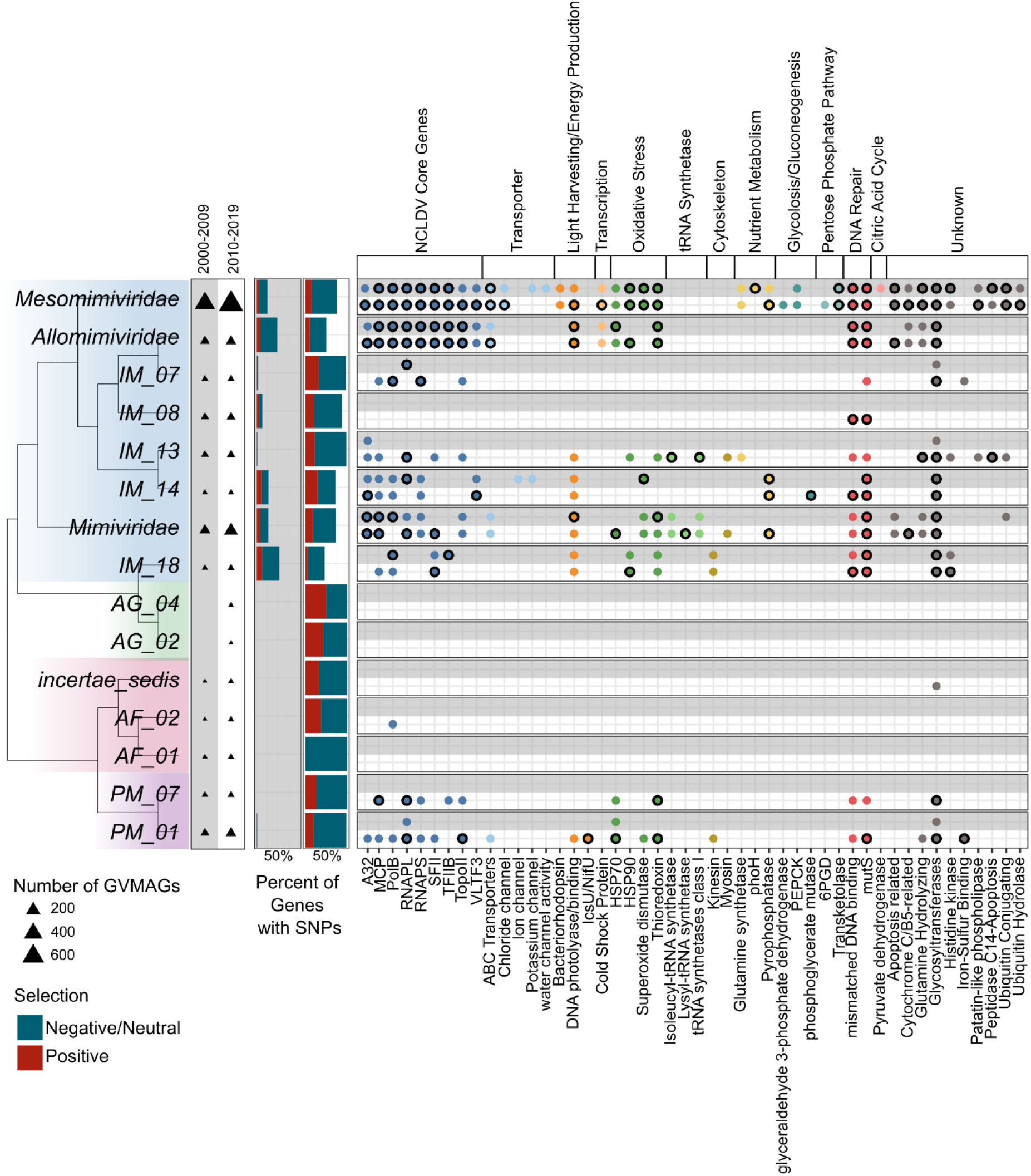
Microdiversity of giant virus families before (2000-2009; shaded grey) and after (2010-2019) spiny water flea invasion. Barplot shows the percentage of genes with single nucleotide polymorphisms (SNPs) that are undergoing positive and negative selection prior and after invasion. Dot plot displays the presence of SNPs in genes of interest. If at least one gene per family and gene category shows positive selection, a black outline was applied. Genes with no dot indicate that the gene does not contain SNPs, it does not denote missing genes. A32: A32-like packaging ATPase; MCP: Major capsid protein; PolB: DNA polymerase family B; RNAPL: RNA polymerase large subunit; RNAPS: DNA-dependant RNA polymerase subunit;SFII: superfamily II helicase; TFIIB: Transcription Initiation factor TFIIB; TopoII: DNA topoisomerase 2; VLTF3: Poxvirus Late Transcription Factor VLTF3 like; PEPCK: phosphoenolpyruvate carboxykinase; 6PGD: 6-phosphogluconate dehydrogenase.

The observed post-invasion increase in SNP rates raises a critical question: did adaptive evolution specifically accelerate following ecosystem disruption? To address this question, we focused specifically on genes showing signatures of positive selection (defined by pN/pS > 1, where pN and pS represent observed nonsynonymous and synonymous polymorphism rates). After implementing the same rigorous controls for sampling effort used in our total SNP analysis, we found that positively selected genes were significantly more abundant in the post-invasion period (p < 0.001; rarefaction mean estimate = -0.43). This pattern persisted after controlling for environmental variables, indicating that the invasion-associated increase in adaptive evolution cannot be explained by coincidental environmental changes. Environmental correlates of positive selection largely mirrored those of overall SNP counts: dissolved inorganic carbon and total organic carbon showed positive associations with genes under positive selection (p = 0.008 and p = 0.002, respectively), while total inorganic carbon and dissolved organic carbon showed negative associations (p = 0.006 and p = 0.027, respectively). The consistent direction but stronger statistical significance of these relationships for positively selected genes (compared to total SNPs) suggests that adaptive evolution responds more sensitively to environmental conditions than overall mutation accumulation. Collectively, these findings provide evidence that the spiny water flea invasion triggered accelerated adaptive evolution in giant virus genomes, potentially reflecting selection for novel host exploitation strategies or altered competitive dynamics in the restructured ecosystem.

While the majority of genes showing evidence of positive selection had no known functional annotation, highlighting the need for further investigation, we did identify selection acting on several giant virus hallmark genes likely involved in host interactions (Fig. 4). The relationship between host interactions and viral evolution became particularly evident when examining specific cases of adaptation following ecosystem disturbance. For example, thioredoxin genes in *Pimascovirales* genomes showed positive selection specifically after the spiny water flea invasion. Thioredoxin likely mediates oxidative stress responses and is actively released by giant viruses during infection (Nishinaka *et al*., 2001; Schrad *et al*., 2020), suggesting adaptation in viral control of cellular redox environments. Similarly, IM_13 genomes showed positive selection in isoleucyl-tRNA synthetase genes following invasion. This enzyme is expressed during infection by the marine giant virus CroV (Cafeteria roenbergensis virus) when infecting zooplankton, and IM_13’s expanded associations with zooplankton hosts after invasion (Fig. 3b) suggest that tRNA synthetase adaptation may have helped to facilitate host range expansion (Yi *et al*., 2024). Quantitatively, we identified significant post-invasion increases in positively selected genes across key functional categories: N-methyltransferase activity (16 to 33 GVMAGs), DNA-dependent RNA polymerase (19 to 29 GVMAGs), major capsid protein (3 to 14 GVMAGs), and glycosyl transferase family 2 (3 to 12 GVMAGs). Additionally, we identified several functional categories showing positive selection exclusively after invasion: induction of catabolism of host mRNA (8 GVMAGs), ubiquitin-conjugating enzyme activity (6 GVMAGs), DNA ligase (NAD+) activity (6 GVMAGs), and ribonucleotide reductase, barrel domain activity (6 GVMAGs). This functional profile of adaptation strongly suggests that giant viruses responded to ecosystem disruption through molecular adaptation in systems critical for host recognition, infection, and replication efficiency.

### Giant viruses occupy an evolutionary middle ground between bacteria and smaller viruses

To place giant virus evolution in a broader evolutionary context, we conducted, to our knowledge, the first systematic comparison of evolutionary rates across three major groups from the same ecosystem: giant viruses, bacteria, and smaller viruses (predominantly Caudoviricetes phages; Supplementary Table S7). Our analysis quantified both whole-genome single nucleotide substitution (SNS) rates and observed nonsynonymous substitutions (obsN) in protein-coding genes (Fig. 5a,b). This approach revealed that giant viruses and bacteria showed statistically indistinguishable whole-genome substitution rates, while smaller viruses evolved significantly faster than both groups. This finding establishes giant viruses as evolutionary intermediates that share genomic stability characteristics with cellular organisms despite their viral nature. The similarity in evolutionary rates between giant viruses and bacteria is particularly interesting given their fundamentally different biological organization and genome sizes. We propose that this convergence in evolutionary rate reflects shared constraints: like bacteria, giant viruses encode complex metabolic and regulatory systems that likely impose similar selective penalties on mutation accumulation. Both bacteria and giant viruses must maintain numerous interdependent functions within a single genomic system, potentially limiting tolerable mutation rates compared to smaller viruses with more streamlined genomes. The significantly higher substitution rates in smaller viruses likely reflect their simplified genetic architecture, which permits greater mutational flexibility while maintaining function.

**Figure 5:**
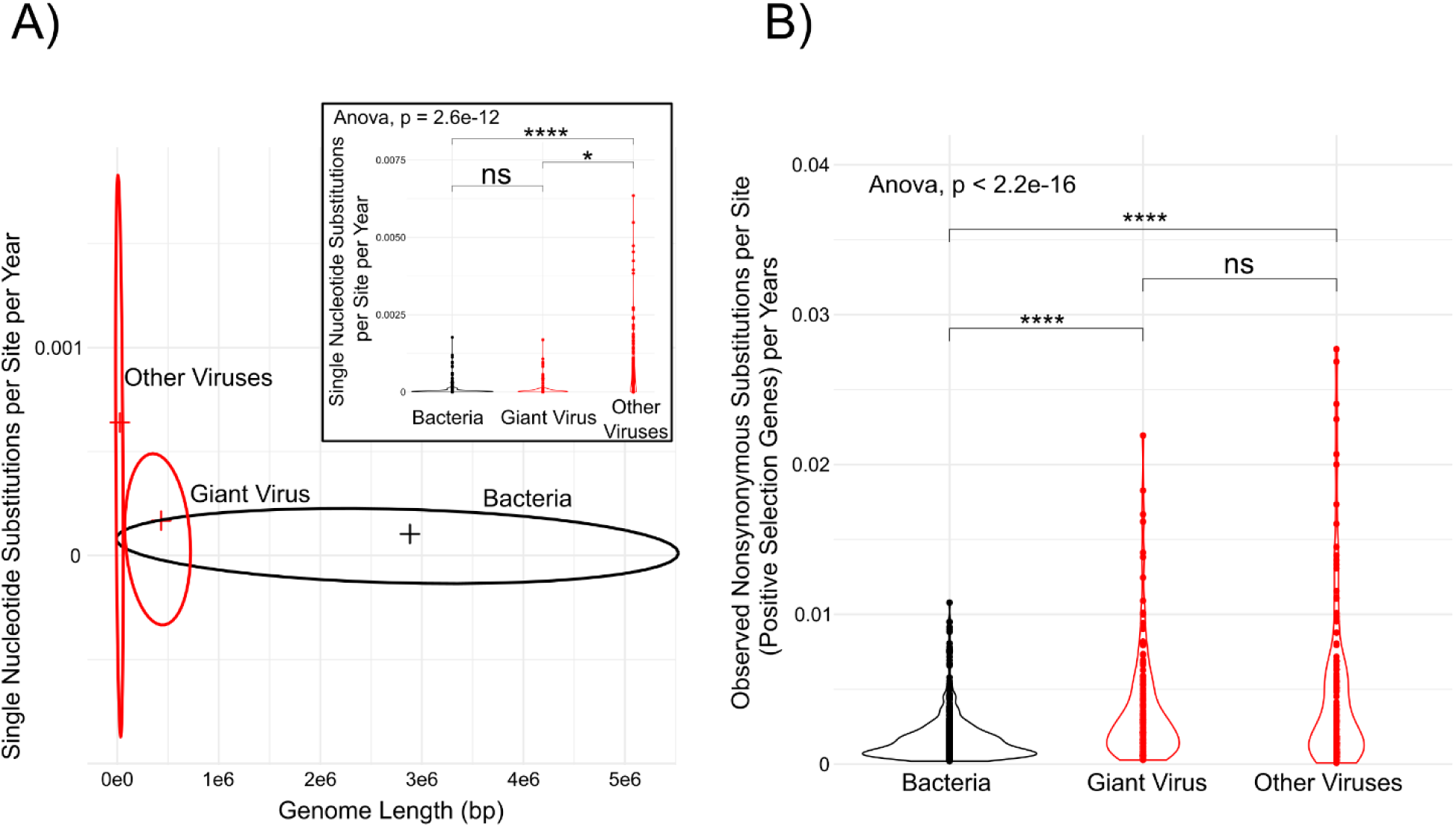
Evolutionary rates of Bacteria, giant viruses and other viruses in Lake Mendota. A) Rates of single nucleotide substitutions by genome length. Inset displays rates as violin plots and Wilcox tests to compare between genomes. B) The number of nonsynonymous substitutions within protein coding genes that exhibit positive selection according to microdiversity analysis. A Wilcox test was used to compare between genomes.

Despite this broad similarity in overall substitution rates, a more nuanced pattern emerged when we focused specifically on genes under positive selection. In these adaptively evolving protein coding genes, giant viruses showed nonsynonymous substitution rates that differed significantly from bacteria but not from other viruses. This finding reveals a fundamental duality in giant virus evolution: while their overall genomic conservation patterns match bacteria, their adaptive evolution dynamics mirror those of other viruses. This hybrid evolutionary character suggests that giant viruses may maintain bacterial-like genomic stability in core functions while retaining virus-like flexibility in genes directly involved in host interactions and environmental adaptation. Previous studies have hypothesized evolutionary rate similarities between giant viruses and cellular organisms (Holmes, 2011; Moelling and Broecker, 2019), but empirical evidence from natural systems has been lacking. Our analysis, leveraging a 20-year time series from a single ecosystem, offers the first direct empirical evidence and takes important steps toward understanding these evolutionary relationships. By demonstrating that giant viruses evolve at rates comparable to co-occurring bacteria while smaller viruses evolve substantially faster, we establish a quantitative framework for understanding giant virus evolution. This positioning of giant viruses as evolutionary intermediates offers insight into both their origins and their ecological success in diverse environments.

## DISCUSSION

Giant viruses are increasingly recognized as important members of microbial communities, yet their evolutionary dynamics in natural settings have remained largely theoretical rather than empirical. While recent metagenomic studies have begun illuminating giant virus diversity and distribution in marine systems, equivalent insights from freshwater ecosystems have been limited to snapshot studies lacking temporal depth. Our reconstruction of 1,512 giant virus genomes from a single lake over two decades transforms this landscape, positioning Lake Mendota as the currently most comprehensively characterized freshwater system for giant virus evolution. This dataset’s unique temporal resolution has enabled us to address fundamental questions about giant virus evolution that were previously restricted to laboratory settings. Our findings reveal that giant viruses employ sophisticated evolutionary strategies to maintain long-term persistence while adapting to ecological disruption.

Our findings both confirm and extend previous observations of temporal patterns in giant virus communities (Roux *et al*., 2017; Gran-Stadniczeñko *et al*., 2019; Laperriere *et al*., 2024; Zhang *et al*., 2024; Fang *et al*., 2025). While earlier studies identified seasonal fluctuations in giant virus abundance, our 20-year time series reveals more complex dynamics: taxonomic groups exhibit distinct persistence patterns, with *Imitervirales* maintaining continuous presence while other orders show intermittent detection across years. This consistent dominance of *Imitervirales*, particularly *Mesomimiviridae*, suggests these viruses possess ecological advantages in freshwater systems, likely stemming from broader host ranges. Additionally, the high gene conservation across our dataset (86% of genes in shared orthogroups) indicates strong functional convergence. This unexpected level of gene sharing likely reflects adaptation to Lake Mendota’s relatively homogeneous physicochemical conditions compared to the pronounced environmental gradients characterizing ocean basins. The combination of persistent dominant lineages and high functional conservation hints that successful giant virus adaptation in freshwater lakes may favor evolutionary stability rather than continuous turnover.

By leveraging our time-series approach, we provide the first direct observation of multiple genomic innovation mechanisms operating simultaneously in giant virus populations. Gene duplication emerged as the predominant driver of genomic expansion (23% of all genes), with functional enrichment in mechanisms used for host infection and viral replication (*e.g.*, ubiquitination pathways). This finding further provides empirical support for recent experimental work suggesting duplication as a primary mechanism of giant virus genome growth (Machado *et al*., 2023). Horizontal gene transfer also proved to be a driver of genome innovation (29% of genes) and showed striking lineage-specific biases, with *Imitervirales* genomes containing ten-fold more transferred genes than *Pandoravirales*. Our temporal resolution allowed us to directly observe 30 gene gain and loss events occurring in real time, largely driven by duplication rather than acquisition from cellular sources. The observation that 16 of 17 gained genes represented duplications of existing viral genes provides further evidence that gene duplication events drive giant virus innovation in natural settings. These real-time observations advance our understanding of the “genomic accordion” model (Filée, 2015), demonstrating that giant viruses simultaneously employ multiple genomic innovation strategies while pruning non-essential functions.

Our putative host association analyses revealed a diverse range of eukaryotic lineages connected to giant viruses in Lake Mendota. The network encompassed well-established giant virus hosts like Stramenopiles alongside numerous putative host relationships spanning Alveolata, Opisthokonta, and other eukaryotic supergroups (Sun *et al*., 2020). Many of these computationally predicted relationships align with recent marine metagenomic studies (Endo *et al*., 2020; Meng *et al*., 2021), suggesting conserved host range patterns across aquatic environments. Importantly, these computational predictions are increasingly validated through independent methods; for instance, the *Mesomimiviridae*-*Ciliophora* relationship identified in our network was recently confirmed through single-cell viral-host interaction studies (Fromm *et al*., 2024). The breadth of potential hosts significantly expands our understanding of giant virus ecology beyond the handful of laboratory-established model systems. Such computational host predictions provide valuable guidance for targeted isolation efforts, potentially addressing the substantial disconnect between metagenomic diversity and cultivated representatives. Beyond their taxonomic implications, these predicted virus-host relationship patterns may serve as sensitive indicators of ecosystem disturbance (Wheatley, Holtappels and Koskella, 2024), as evidenced by the emergence of multiple new associations with dinoflagellates following the spiny water flea invasion. The ability to identify such ecological reorganizations through virus-host network analysis demonstrates the potential of viral ecology as a tool for monitoring freshwater ecosystem health and resilience.

The 2009 spiny water flea invasion created an unprecedented natural experiment to investigate viral adaptation to ecosystem disruption. Our statistical analyses, controlling for sampling effects, environmental parameters, and temporal autocorrelation, revealed that giant viruses responded to this perturbation through a dual strategy: retention of established host relationships coupled with opportunistic expansion into newly available host niches. Post-invasion networks showed significant increases in both the number and stability of predicted virus-host associations, with new connections forming with taxa that increased in abundance following invasion. Distance-based redundancy analysis confirmed that invasion status was the dominant explanatory variable for community restructuring. This ecological response was mirrored at the genomic level, where single nucleotide polymorphism analyses revealed an increase in microdiversity following invasion and an increase in the proportion of genes under positive selection. The functional profile of positively selected genes suggests targeted adaptation of genes involved in host exploitation. Different viral families showed different intensities of selection, with some showing minimal change while others experiencing increased adaptive evolution, highlighting the complex selective landscape even within a single ecosystem. The evolutionary flexibility of giant viruses likely enhances long-term persistence in a fluctuating environment, allowing exploitation of newly available host niches without sacrificing established infection networks.

Our comparative evolutionary rate analyses positions giant viruses as evolutionary intermediates between cellular life and other viruses (*e.g.,* bacteriophages). Whole-genome substitution rates in giant viruses matched those of co-occurring bacteria, yet lagged the accelerated rates of smaller phages from the same ecosystem. This finding suggests that genome size and complexity, not cellular structure, may be a primary determinant of evolutionary tempo. Intriguingly, when focusing specifically on adaptive evolution in positively selected genes, giant viruses showed nonsynonymous substitution patterns statistically similar to smaller viruses but significantly different from bacteria. This creates a fascinating evolutionary duality, giant viruses maintain bacterial-like genomic stability in core functions while retaining virus-like adaptive flexibility in genes directly involved in environmental response. Our findings thus offer insight into not only how giant viruses evolve, but also increases the support for their ambiguous position in the biological world, neither fully virus-like nor bacteria-like in their fundamental evolutionary behavior.

The two-decade record from Lake Mendota increases our understanding of giant virus evolution, demonstrating that viruses are capable of sophisticated evolutionary responses to environmental change. Our findings reveal a multi-faceted adaptive strategy: continuous gene content innovation through duplication, horizontal gene transfer, and occasional de novo emergence; flexible host range expansion rather than replacement following ecosystem disruption; and a hybrid evolutionary mode that combines bacterial-like genomic stability with virus-like adaptive capacity. This evolutionary versatility likely explains how giant viruses have achieved ecological dominance across diverse aquatic environments. Beyond advancing viral evolution theory, these findings have significant implications for ecosystem modeling, as they demonstrate that viral communities respond to disturbance in ways that likely stabilize rather than amplify ecosystem perturbations. While our study provides the first comprehensive, vicennial analysis of giant virus evolution in any ecosystem, important questions remain. Future work should employ single-virus genomics and multiomics to validate host predictions and determine the expression patterns of putatively adaptive genes. Comparative time-series analyses across multiple lakes with different disturbance histories will reveal whether the evolutionary patterns observed here represent universal strategies or lake-specific adaptations. Ultimately, integrating viral evolutionary dynamics into ecosystem models will be essential for understanding and predicting microbial community responses to environmental change.

## METHODS

### TYMEFLIES Dataset

471 water filter samples were collected from the eutrophic freshwater Lake Mendota in Madison, WI, USA. Samples span the years 2000 - 2019 and include multiple samples from different microbial seasons (*i.e.*, Ice-on, Spring, Clearwater, Early Summer, Late Summer and Fall; defined by Rohwer *et al*., 2023). The collection of these sequences are designated as the Twenty Years of Metagenomes Exploring Freshwater Lake Interannual Eco/evo Shifts (TYMEFLIES) dataset (Rohwer and McMahon, 2022; Rohwer *et al*., 2025). Co-assembly with MetaHipMer performed on the TYMEFLIES dataset (Oliver *et al*., 2024) resulted in 1,535 possible GVMAGs. Average Nucleotide Identity (ANI) calculated using fastANI v.1.34 (Jain *et al*., 2018) did not result in any removal of genomes when based at 96%. GVMAGs were then subjected to taxonomic classification using GVClass v.1.0 (Pitot, Brůna and Schulz, 2024). MAGs that were not predicted as *Nucleocytoviricota*, or contained a high number of contamination with GVClass were removed from further analysis (n = 23), leaving 1,512 non-redundant GVMAGs for further analysis. For family-level analyses, only genomes with inferred family taxonomy were retained. Pandoravirales was excluded from family-level analyses as there was no family-level taxonomic classification identified by GVClass at the time of publication. To calculate relative abundance of GVMAGs, mapping was performed with bowtie2 v.2.5.3 (Langmead and Salzberg, 2012) and converted to bam with samtools v.1.19.2 (Li *et al*., 2009).

Using inStrain v.1.9.0 (Olm *et al*., 2021), a GVMAG within a given sample was considered present if a minimum of 20% of the viral genome was covered by reads with an average depth of at least 1. This allowed us to visualize the unique number of GVMAGs per year and the absolute number of GVMAGs per year. To calculate abundance of GVMAGs per season, we used the bam output and raw read counts were aggregated at the family level. Only samples present with collection metadata (*i.e.,* samples with Year, Month and Day recorded) were retained for analysis. Differential abundance analysis was performed using ANCOM-BC2 (Analysis of Compositions of Microbiomes with Bias Correction 2) (Lin and Peddada, 2024), which accounts for the compositional nature of microbiome data by modeling log-ratios of abundances rather than raw counts. ANCOM-BC2 tests for differential abundance of taxa while controlling for compositionality bias inherent in relative abundance data. The implementation included a fixed effect for seasonal periods to identify taxa that significantly change across seasons, and a random effect for collection year to account for temporal pseudoreplication in the sampling design. The ANCOM-BC2 analysis incorporated several critical parameters: a structural zero detection mechanism with a prevalence threshold of 10% (prv_cut = 0.10) to distinguish between true zeros and sampling zeros; a minimum library size filter (lib_cut = 1000) to remove low-quality samples; and a small effect size threshold (s0_perc = 0.05) to improve sensitivity. Multiple hypothesis testing correction was applied using Holm’s method to control family-wise error rate. Four complementary statistical tests were performed: (1) global tests to identify taxa that show any significant differences across all seasons, (2) pairwise tests to detect differences between specific season pairs, (3) Dunnett’s tests to compare each season against a reference season, and (4) trend tests to identify taxa showing monotonic patterns across the seasonal gradient. This comprehensive testing approach enabled detection of various patterns of temporal dynamics in microbial families. The resulting bias-corrected abundances were log10-transformed and averaged across samples within each seasonal category to generate a heatmap visualization matrix of family-level abundance patterns throughout the seasonal cycle.

### Phylogenetic Tree Building

Phylogenetic trees of GVMAGs were reconstructed using nsgtree v.0.5 (https://github.com/NeLLi-team/nsgtree), which leverages MAFFT v.7.310 (Katoh *et al*., 2002), IQ-TREE v.2.3.3 (Minh *et al*., 2020) and GVOG7 hidden Markov models (HMMs). Tree visualization was carried out using ggtree in R (Yu *et al*., 2017).

### Eukaryotic host prediction

To assess host-giant virus co-occurrences, we used PolB as a marker gene for *Nucleocytoviricota* and the 18S rRNA gene as a marker gene for eukaryotes. For *Nucleocytoviricota*, we used HMMER and hmmsearch (-E 1e-5) (Finn, Clements and Eddy, 2011) with HMMs generated by Endo *et al*., 2020. Using the entire TYMEFLIES dataset, we recovered 194,957 PolB sequences, although this collection included sequences other than *Nucleocytoviricota* across 471 samples. The number of PolBs per sample ranged from 42 to 1342. We identified 61,426 *Nucleocytoviricota* PolBs using the MAFFT ’addfragments’ option (Katoh *et al*., 2002) and pplacer (Matsen, Kodner and Armbrust, 2010) with the alignment and reference tree provided by Endo *et al*., 2020. Samples with less than 50 sum of raw abundances were removed to avoid low sequencing bias.

For the 18S rRNA gene, we used Infernal and the cmsearch program (Nawrocki and Eddy, 2013) to search the contig files for sequences matching to each of the following small subunit rRNA (SSU rRNA) models from the Rfam database (Kalvari *et al*., 2018): eukaryotic 18S rRNA (RF01960), microsporidial 18S rRNA gene (RF02542), archaeal 16S rRNA gene (RF01959) and bacterial 16S rRNA gene (RF00177). In order to include long SSU rRNA sequences in the analysis, the --anytrunc option in cmsearch was used and the output parsed to find contigs in which sequential and multiple non overlapping alignments to the covariance model were found. We searched the retrieved SSU rRNA sequences against a database that contained all SSU rRNA sequences from both SILVA (Quast *et al*., 2013) and PR^2^ (Guillou *et al*., 2013) databases using BLAST+ (Camacho *et al*., 2009). This workflow is provided here at https://github.com/NeLLi-team/ssuextract. Sequences with no hits to the database were added to a reference SSU rRNA tree to confirm their phylogenetic affiliation at the domain level.The reference tree was built as follows: SSU rRNA from the SILVA and PR^2^ databases were clustered at 75% identity using Usearch (Edgar, 2010) and the resulting centroids were aligned using L-INS-i in MAFFT (Katoh *et al*., 2002). Tree building was done with IQ-TREE (Minh *et al*., 2020) with default parameters including the substitution model selection feature (Kalyaanamoorthy *et al*., 2017). Sequences that branched within a cluster that belonged to a different domain were considered misannotations. We used identity-based sequence clustering at 97% and then at 85% to get operational taxonomic units of the SSU sequences, which had a blast best hit with 18S rRNA gene sequences and those with no blast hit but that branched within the 18SrRNA gene region of the reference tree, together with the sequences from SILVA, PR^2^, SSU gene sequences from eukaryotic genomes from NCBI, and long 18S rRNA gene amplicons from environmental samples (Jamy *et al*., 2022). After discarding singleton clusters we aligned the OTU representative 18S rRNA gene sequences concatenated with their respective 28S rRNA gene sequence, whenever they were extracted from the same contig, using the cmalign program with the RF01960 and RF02543 covariance models, respectively. Columns with more than 90% missing data were removed with trimAl (Capella-Gutiérrez, Silla-Martínez and Gabaldón, 2009). We then built a tree using the IQ-TREE program with the GTR+F+R10 model and 1,000 ultrafast bootstrap replicates using the aligned centroid sequences. We manually inspected the tree and built it again after discarding those sequences with terminal branches with length > 0.5 and those that branched within clades dominated by sequences classified into a different eukaryotic supergroup. Finally, all environmental SSU OTU representative sequences were placed into the tree using the EPA-ng program (Barbera *et al*., 2019) and then taxonomy was assigned with the gappa software (Czech, Barbera and Stamatakis, 2020) using the PR^2^ taxonomy.

Using similar methods from Meng *et al*., 2021, we created a PolB and 18S rRNA gene matrix to perform network inference. We first applied a centered log-ratio (clr) transformation on each matrix separately, after adding a pseudo count of one to all values. We also filtered out low-abundance genes using a lower quartile filtering approach. Finally, we used a combined matrix as the input for Flashweave v.0.19.2 (sensitive=true, heterogeneous=false,alpha = 0.01) (Tackmann, Rodrigues and Mering, 2019). Following results from Flashweave, we recreated two *Nucleocytoviricota* PolB phylogenetic trees. The first to be used to classify PolBs at the family level, we used the GVOGm0054 marker and aligned PolB sequences with reference sequences using MAFFT and FastTree (Aylward *et al*., 2021). Any PolBs not within our families of interest (*i.e.*, those included in Figure X) were removed from visualization. The second tree was used to confirm relationships using the Taxon Interaction Mapper (TIM) (Kaneko *et al*., 2021), however, these only included sequences with at least 500 amino acid length resulting in 3,137 polB sequences.

To estimate the strength of associations from the Flashweave output, we calculated an association score. First, we clustered *Nucleocytoviricota* PolB sequences using CD-HIT v.4.8.1 (Fu *et al*., 2012), then used this clustering group to find consecutive associations and the max edge per *Nucleocytoviricota*-eukaryote pair. We only kept clusters that had at least three genomes. Our formula includes a decay factor using a decay rate from Hewson *et al*., 2012 for a Phycodnaviridae virus converted into decay per year. Additionally, we applied a small bonus factor of 1.1 for associations that were seen in consecutive years. We then took the max edge value at each year into consideration. The resulting formula:

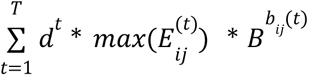

where *d* is the decay constant, 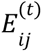 is the edge between an *Nucleocytoviricota* and eukaryote in that year, and 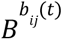 is the bonus factor to the power of the number of current consecutive associations in that year. Using this weighted score, we found the median per *Nucleocytoviricota*-eukaryote pair (removing any pairs that only had one cluster).

Finally, we visualized changes in associations in the Flashweave network output before and after the spiny water flea invasion in Lake Mendota. We implemented multiple analytical approaches to empirically determine when significant network restructuring occurred. First, we calculated annual network metrics after preprocessing our data consistently by aggregating edges to their maximum weights between each virus-host pair. To objectively identify potential abrupt shifts in network structure, we implemented a changepoint detection analysis using the Pruned Exact Linear Time (PELT) algorithm. This approach analyzes time series data to identify statistically significant points where the underlying distribution of the data changes, particularly in mean or variance. For each network metric (edges, density, and modularity), we constructed annual time series (2000-2018) and applied the PELT algorithm with a penalty parameter determined by the Bayesian Information Criterion to prevent overfitting. We removed 2019 from our analysis as it did not comprise a full calendar year of data. Our analysis did not detect statistically significant changepoints in any of the network metrics, suggesting that network restructuring occurred as a continuous process rather than through discrete shifts detectable at annual resolution.

We then performed a sliding window analysis testing alternative division points between 2006-2012. For each potential breakpoint year, we divided our dataset into ’before’ and ’after’ periods and calculated the absolute difference in network density between periods after aggregating edges by maximum weight between each virus-host pair. To assess statistical significance, we implemented a permutation test (500 permutations) for each breakpoint. In each permutation, we randomly reassigned samples to ’before’ and ’after’ periods while maintaining the original sample sizes, recalculated network metrics, and computed the density difference. The p-value for each breakpoint represents the proportion of permutations that produced a density difference greater than or equal to the observed difference. This approach tests whether the observed network differences are statistically distinguishable from those arising by random temporal partitioning of the data. None of the tested breakpoints yielded statistically significant results, with p-values ranging from 0.162 (2009) to 0.456 (2011). This suggests that while visual inspection of network metrics indicates changes in community structure around the time of spiny water flea detection, these changes were not statistically significant to be distinguished from the background of temporal variability when analyzed at annual resolution. We caution that our network plot should be interpreted as contrasting two distinct ecological states: a pre-established and post-established invasion system.

To assess the robustness of observed virus-host association changes while accounting for temporal differences in sampling intensity in the pre-invasion (2000-2009) and post-invasion (2010-2019) periods, we implemented a mixed-effects logistic model that included yearly sequencing effort as a covariate confirmed that the probability that any virus–host pair formed an edge remained significantly higher in the post-invasion period (Edge_presence ∼ Period(pre/post) + log(Samples) + (1|Year)). We then used a bootstrapping approach with 1,000 iterations to randomly choose samples from the period with the greater number of samples (*i.e.*, post). For each iteration, we randomly sampled (with replacement) an equal number of samples from both time periods, constructed the network, and recorded which virus-host pairs were present. This generated a frequency distribution of associations, allowing us to calculate the proportion of bootstrap networks in which each association appeared. We considered associations appearing in ≥95% of bootstrap networks as high-confidence associations. This approach permitted statistical discrimination between robust ecological patterns and potential sampling artifacts, particularly for identifying associations that consistently appeared or disappeared between periods despite sampling variation.

Additional network analyses were performed to ensure the robustness and statistical validity of observed shifts in virus-host associations. A bias-aware null model was used to test whether observed virus-host associations were significantly enriched or depleted compared to random expectations, we generated 5,000 degree-preserving random graphs and calculated Z-scores and empirical p-values for each virus-host subdivision combination. A down-sampling (rarefraction) analysis was performed to account for sampling bias due to increased sequencing depth post-invasion. We randomly subsampled the post-invasion network to match the pre-invasion edge count, repeating this process 5,000 times to ensure robustness of observed patterns. A leave-top-hubs-out robustness analysis was performed by systematically removing highly connected viral nodes to test network stability and assess whether observed structural patterns were robust to the loss of key taxa. Finally, a permutation test on year by shuffling labels of network edges to statistically confirm that structural changes in the network were not due to random temporal variation. Detailed descriptions of these additional analyses can be found in the Supplementary Methods.

### Identification of Gene Duplication & HGTs

To identify gene duplications, we used Orthofinder v.2.5.5 (Emms and Kelly, 2019) to find orthogroups and putative gene duplications in giant virus genomes, resulting in 29,649 orthogroups. For horizontal gene transfer (HGT) identification we used BLASTp (settings: --evalue 1e-5 --id 30 --max-target-seqs 50) with diamond using a custom-made database from Lake Mendota bacterial single assembly bins as well as the NR database (Buchfink, Reuter and Drost, 2021). We identified HGTs as those that contained a percent of identical positions (pident) as 50%. Putative HGTs above 80% were labeled as recent HGTs as this most likely indicated a recently acquired viral gene. Finally, we considered putative de-novo genes as those that belonged to a single orthogroup (*i.e*., ORFan) and had no homolog to the NR database or Lake Mendota single assembly bacterial bins. It is important to note that these de-novo genes could have been transferred from other species not in this set of genomes, thus creating artifacts of ORFans. To annotate genes, we used eggNOG-mapper v.2.1.12 (Cantalapiedra *et al*., 2021) with database v.5.0.2 and pfam_scan v.1.6 with a database from Nov 2019.

Using results from inStrain, we considered genes as present if they contained at least one coverage and 50% breadth (Olm *et al*., 2021). For every gene considered at absence in a sample, we checked if the surrounding genes were considered present. This was applied in order to exclude possible gene candidates that contained multiple genes on the same contig in close proximity not mapped due to insufficient mapping rather than actual gain/loss.

Furthermore, we removed any genes left that contained higher mapping (>1) or those with high breadth (>50%) in the dataset prior to further analysis. We then used the years the respective giant virus was present in the data and the years the gene was present to determine if the gene could be a candidate for gene gain or gene loss. This method is very conservative as it only considered gene candidates if it passed multiple parameters. For gene gain we only considered genes that occurred at least 3 years after the first presence of the giant virus, had mapping on either side, and persisted until at least two years before the last sampling of the giant virus. For gene loss, we applied a similar process but for genes that occurred at least 3 years before the last presence of the giant virus, had mapping on either side, and persisted at least two years after the first sampling. For each putative gene gain and loss, mappings were manually checked to confirm true gain and loss.

### Evolutionary rates

The bam files generated from the presence/absence method above were further used in Metapop v.0.0.60 for analysis of micro-diversity (settings: –id_min 93 –snp_scale both --min_dep 5 --trunc 5) (Gregory *et al*., 2022). Prodigal genes generated from GVClass v.1.0 (Pitot, Brůna and Schulz, 2024) were used in place of self-annotation by MetaPop (Gregory *et al*., 2022).

To consider the rates of substitution between bacterial, viral (primarily bacteriophages) and giant virus genomes, we used Instrain v.1.9.0 with default settings for profile (Olm *et al*., 2021) and MetaPop for rates within coding sequences. Bacterial and viral genomes included were recovered following methods from Oliver *et al.,* 2024. In our whole genome single nucleotide substitution comparison, we only considered genomes that contained at least a coverage of 5 in each sample and were present in at least 3 years. First, we divided the number of nucleotide substitutions by the length of the genome. We then calculated the mean of the previous value per genome and divided by the year difference between the first year and the last year of observed substitutions. We further filtered to remove genomes that did not have a difference of at least three years between the first year and the last year. In order to find rates within coding genes, we used prodigal-gv in geNomad to annotate other virus genomes (Camargo *et al*., 2024) and Prodigal v2.6.3 (Hyatt *et al*., 2010) to annotate bacteria. For substitutions within genes, we only considered genomes that were present in at least three years, with at least three year difference between the first year and the last year and genes with a pNpS ratio above 1, as considering all genes was consistent with the inStrain results. We took the mean of the number of observed nonsynonymous substitutions divided by the gene length of each genome, and further divided this by the number of years between the first year and the last year of the genome.

## DATA AVAILABILITY

Giant virus co-assembly metagenomes are available under taxon identifier 3300059473 in JGI’s IMG/M platform (https://img.jgi.doe.gov/cgi-bin/m/main.cgi?section=TaxonDetail&page=taxonDetail&taxon_oid=3300059473). Information on single metagenomic studies can be found using JGI’s GOLD database (https://gold.jgi.doe.gov/study?id=Gs0136121). Other data used in this study include: files used for recruiting *Nucleocytoviricota* PolB genes, MAFFT alignment file and reference tree (ftp://ftp.genome.jp/pub/db/community/tara/Biogeography/), environmental data from North Temperate Lakes Long-Term Ecological Research (https://lter.limnology.wisc.edu/all-data/).

## ACKNOWLEDGEMENTS

The work (proposal: 10.46936/10.25585/60001198) conducted by the U.S. Department of Energy Joint Genome Institute (https://ror.org/04xm1d337), a DOE Office of Science User Facility, is supported by the Office of Science of the U.S. Department of Energy operated under Contract No. DE-AC02-05CH11231. We thank Dr. A. Ali Heydari for his valuable input in the association score mathematical formula.

## AUTHOR INFORMATION

United States Department of Energy Joint Genome Institute, Berkeley, CA, USA

Yumary M. Vasquez, Miguel F. Romero, Robert Bowers, Tanja Woyke, Frederik Schulz

Department of Bacteriology, University of Wisconsin at Madison, Madison, WI, USA Katherine McMahon

Department of Integrative Biology, University of Texas at Austin, Austin, TX, USA Robin R. Rohwer

