## Supplemental Information for "Vicennial metagenomic time series unveils evolutionary dynamics of giant viruses in a freshwater ecosystem"

### **Supplementary Figures**

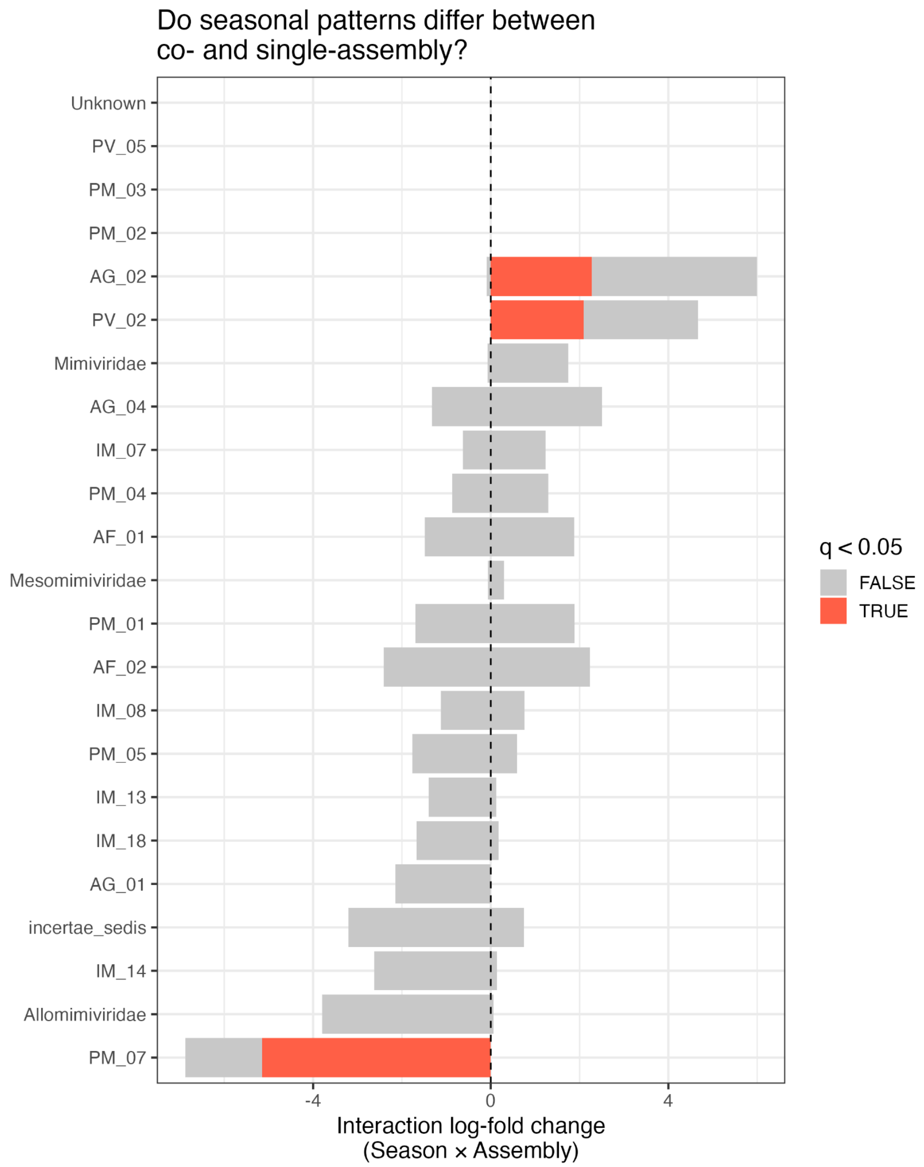

Supplementary Figure S1: Differential seasonal patterns between co-assembly and single-assembly approaches across viral taxa.
Interaction log-fold changes representing how seasonal patterns differ between co-assembly and single-assembly methods across various viral families. The x-axis shows the magnitude and direction of the Season × Assembly interaction, with positive values indicating stronger seasonal effects in one assembly type compared to the other. Each horizontal bar represents a viral family with its corresponding confidence interval. Orange bars indicate statistically significant interactions ($q < 0.05$), while gray bars represent non-significant differences. Only three groups (AG_02, PV_02, and PM_07) demonstrate significant differences in seasonal patterns between assembly methods, with PM_07 showing a negative interaction effect. The vertical dashed line at x=0 represents no interaction between seasonal patterns and assembly method.

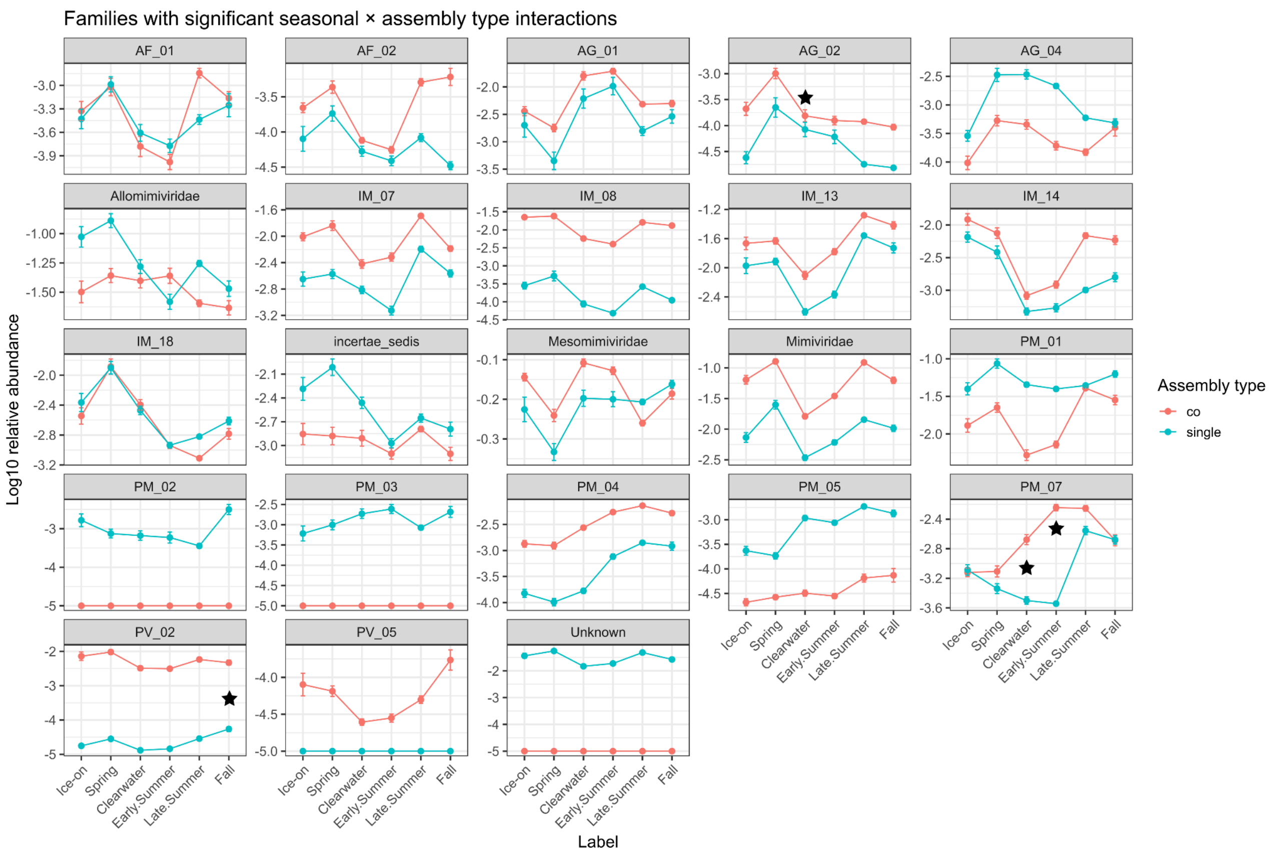

Supplementary Figure S2: Seasonal abundance patterns of viral families showing significant interactions between season and assembly methodology.
Plot displays the log10 relative abundance of viral families across six seasonal time points using two different assembly approaches. Black stars highlight particularly significant interaction points in specific families (AG_02, PM_07, and PV_02). We did not find any genomes for PM_02 or PM_03 in the co-assembly. All Average Nucleotide Identity (ANI) clusters for these species are binned with single assembly sequences only.

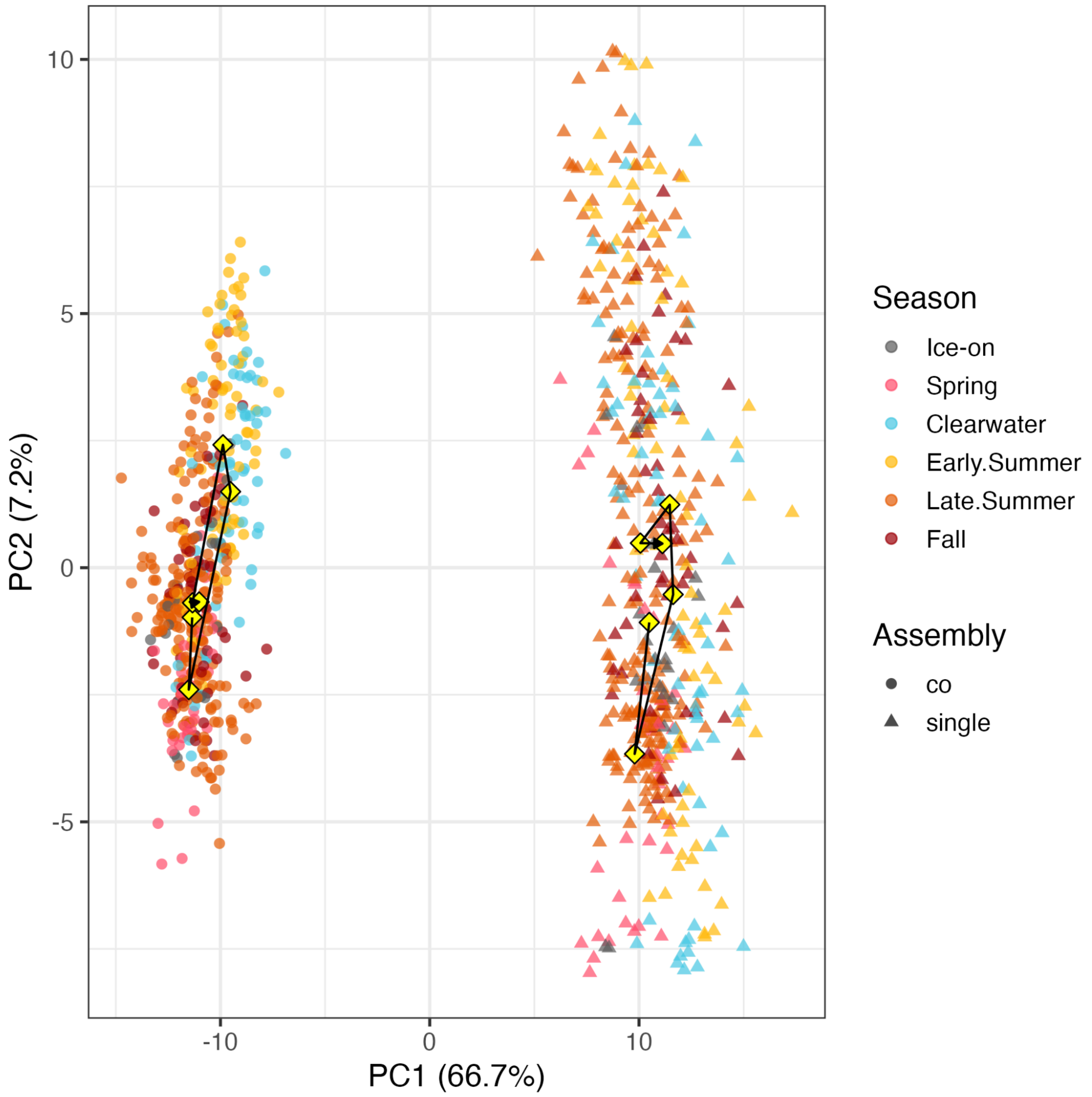

Supplementary Figure S3: Principal component analysis (PCA) of viral community composition between the two methodological differences.
Co-assembly samples (circles) clustering on the left and single-assembly samples (triangles) on the right. PC2 (7.2% variance) captures seasonal patterns, with samples colored by season (Ice-on, Spring, Clearwater, Early Summer, Late Summer, and Fall). Black lines connect seasonal centroids (yellow diamonds) within each assembly method, showing parallel seasonal trajectories despite the methodological separation.

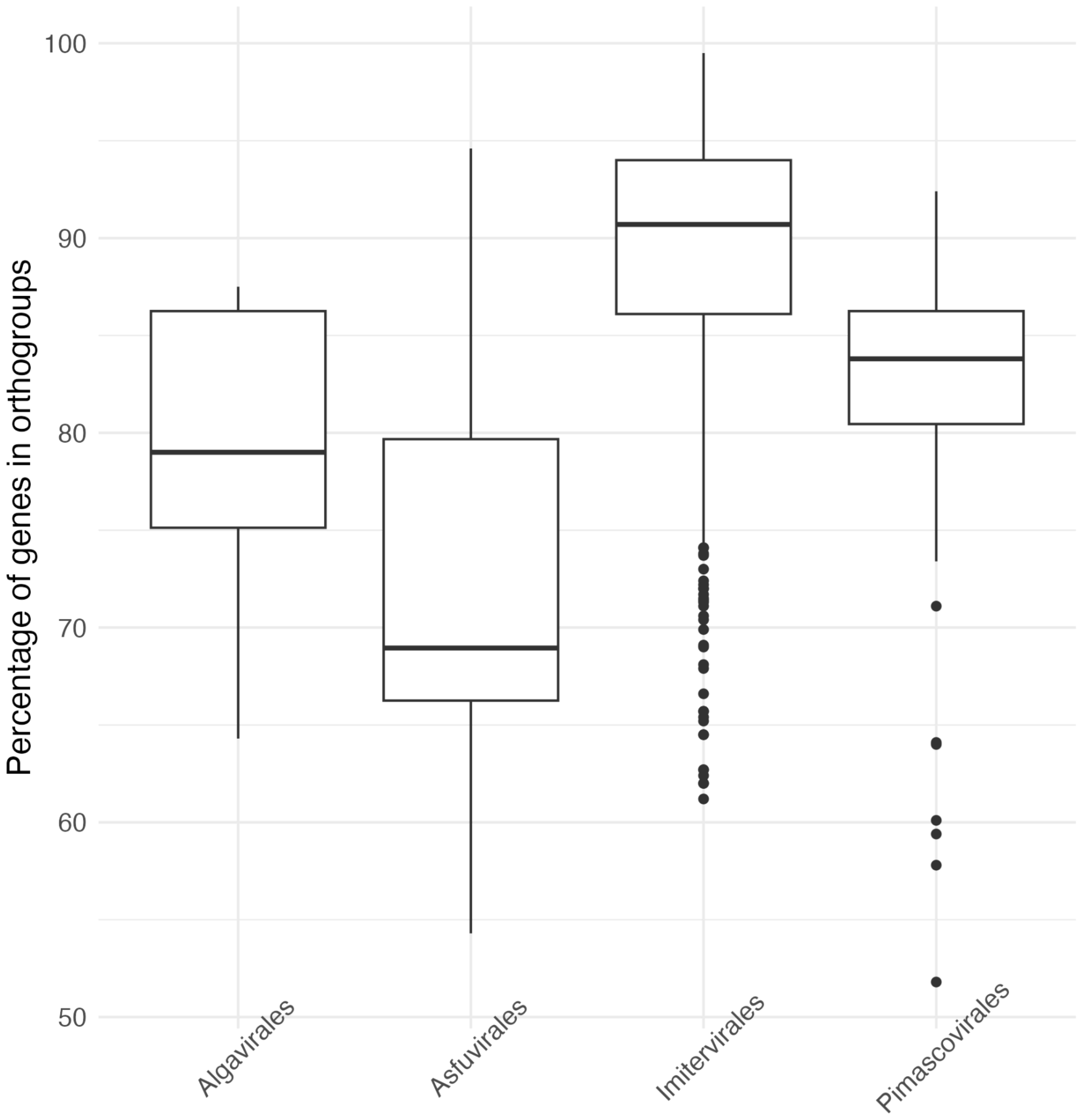

Supplementary Figure S4: Comparison of orthologous gene content across viral orders.
Box plots display the percentage of genes assigned to orthogroups for Algavirales, Astuvirales, Imitervirales, and Pimascovirales calculated on a per-genome basis. Imitervirales, which had the highest number of recovered GVMAGs, showed the highest proportion of genes assigned to orthogroups.

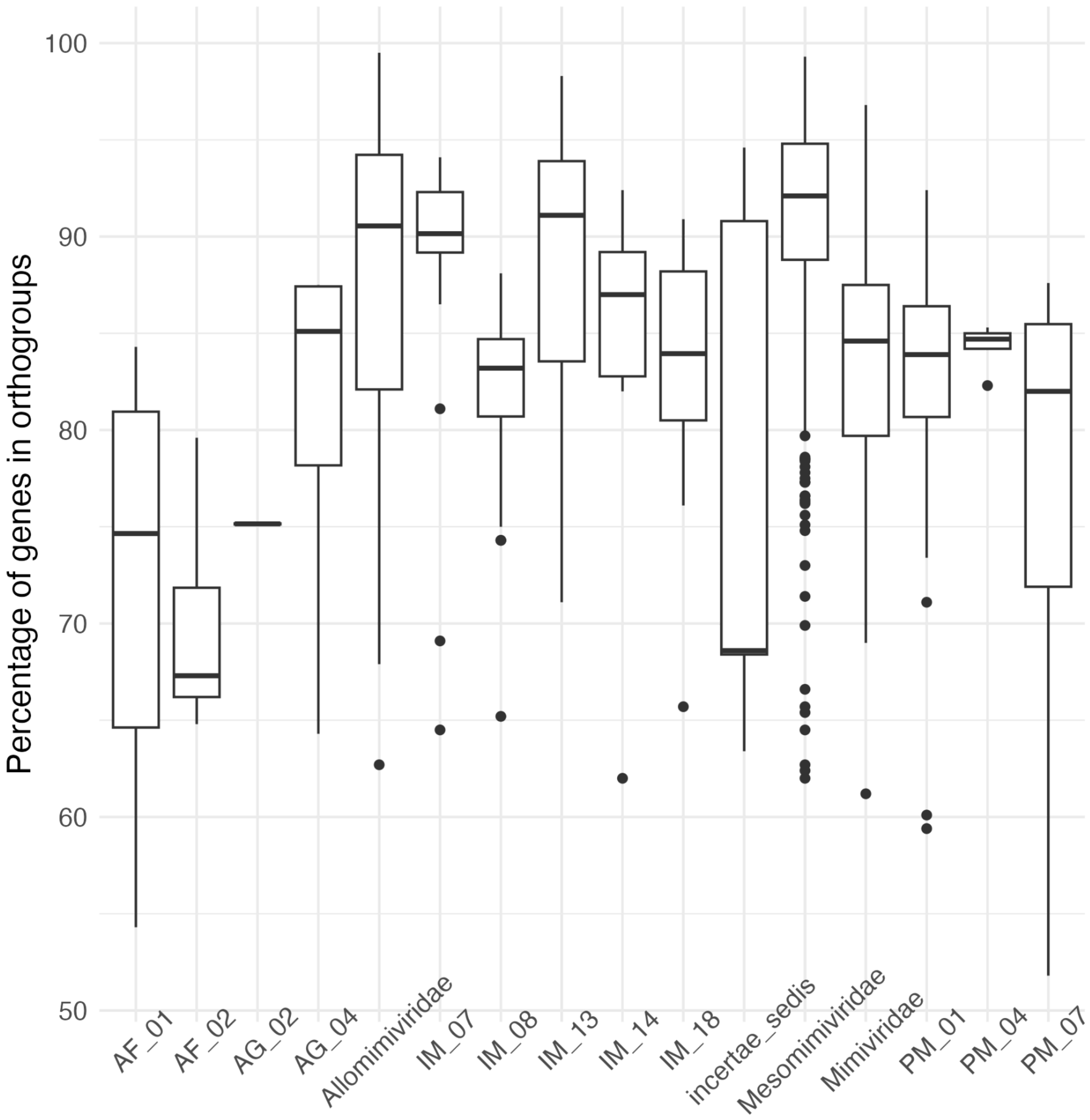

Supplementary Figure S5: Comparison of orthologous gene content across viral families. Box plots display the percentage of genes assigned to orthogroups for multiple giant virus families calculated on a per-genome basis.

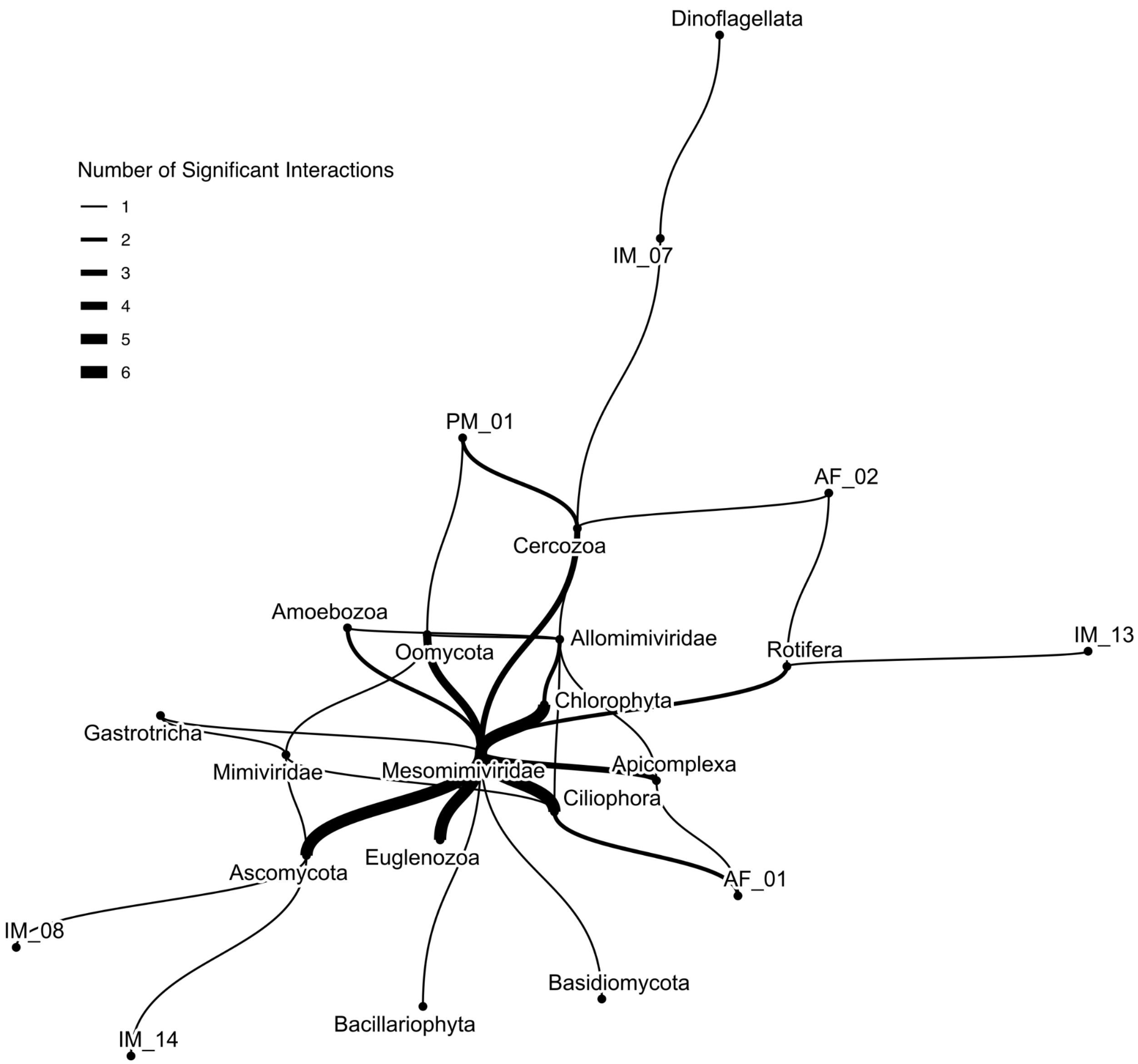

Supplementary Figure S6: Network of significant co-occurrence relationships using the Taxon Interaction Mapper (TIM) (Kaneko et al., 2021).
The thickness of connecting lines represents the number of significant interactions detected between nodes. Mesomimiviridae appears as a central hub with numerous strong connections to eukaryotic groups. Other viral groups display more specialized relationships, such as IM_07 connecting primarily to Dinoflagellata. Sequences of least 500 amino acid length were included (3,137 polB sequences).

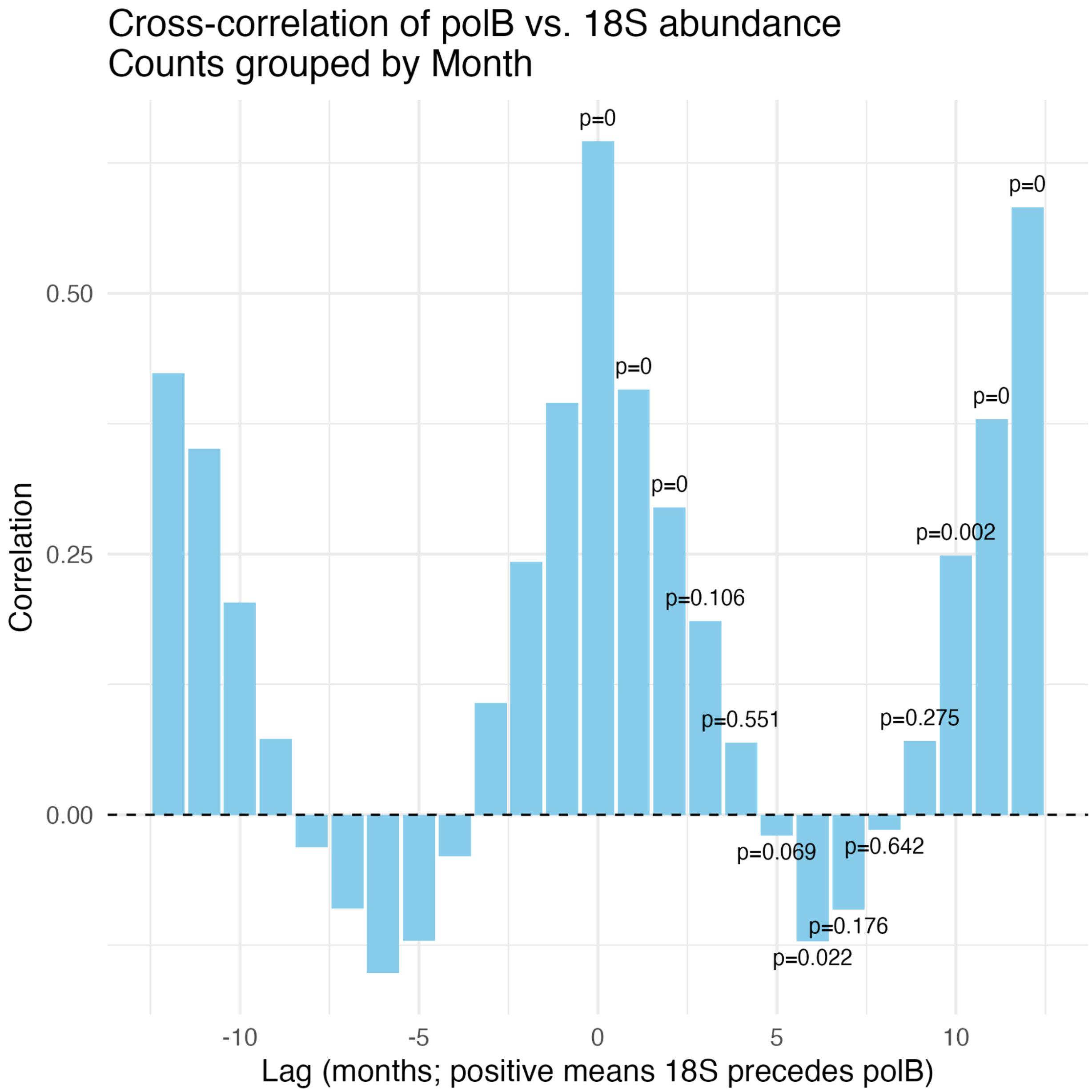

Supplementary Figure S7: Cross-correlation analysis (CCF) of monthly viral polB abundance versus host 18S rRNA gene abundance.

The vertical bars represent the strength of correlation (cross-correlation coefficient, y-axis) at different monthly lags (x-axis). Positive lags indicate that host 18S rRNA gene abundance precedes viral polB abundance, while negative lags indicate the opposite. Labels indicate p-values from lagged linear regression models at specific lags. The strongest and most significant correlation occurs at lag 0 (same-month correlation), suggesting immediate coupling between host abundance and viral abundance. Smaller significant correlations at positive lags (e.g., lag 1 and lag 2 months) indicate potential delayed effects, suggesting that increases in host abundance may slightly precede increases in viral abundance.

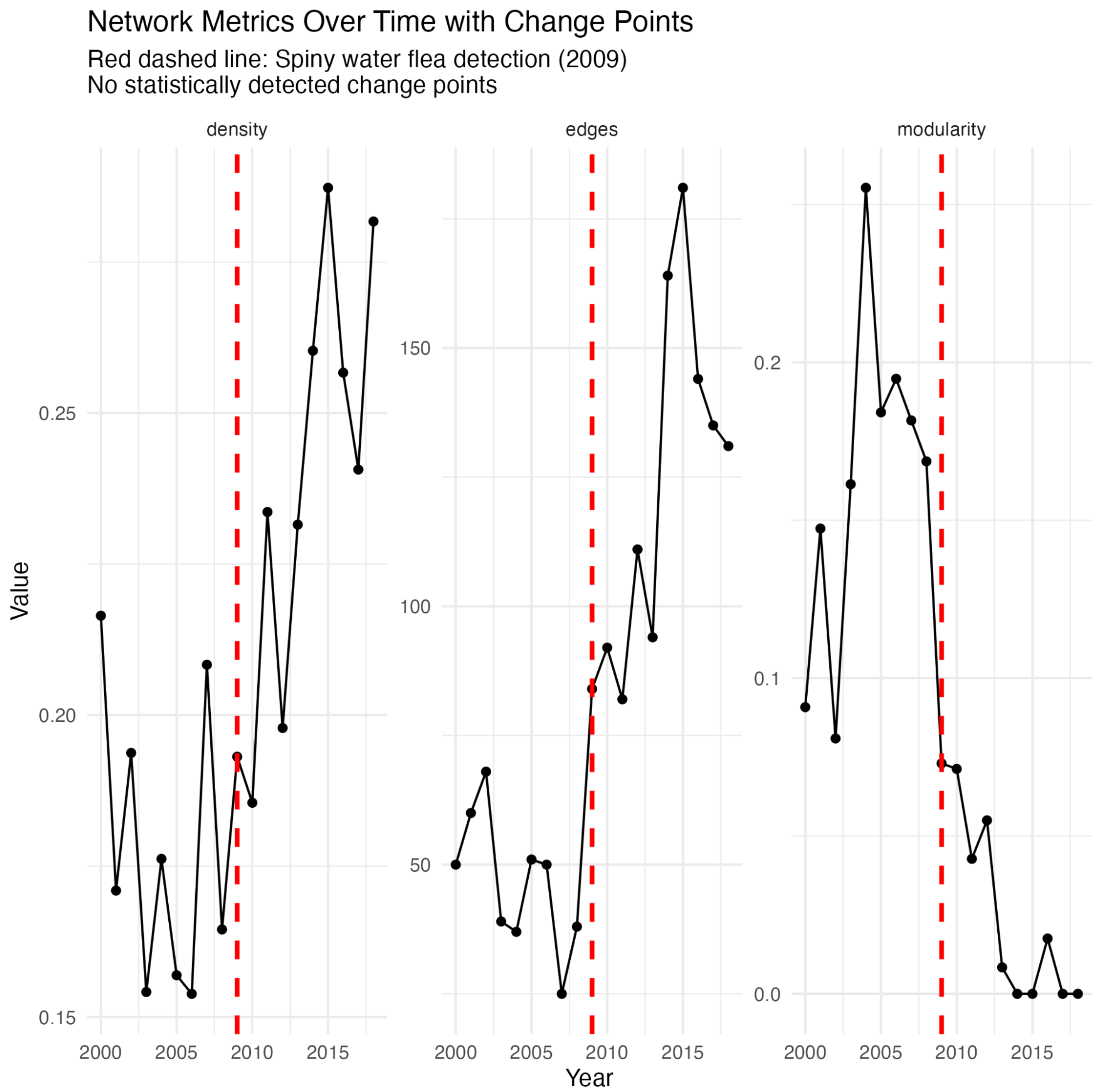

Supplementary Figure S8: Temporal evolution of key network metrics characterizing the Nucleocytoviricota-eukaryote interaction networks in Lake Mendota from 2000-2018. Panels show: (A) Network density (proportion of potential connections that are actually realized), which peaked in 2010 rather than 2009, revealing that network interconnectedness intensified following initial invasion; (B) Total edges (number of distinct virus-host associations), which increased sharply in 2009 coinciding with spiny water flea detection, indicating immediate formation of new interaction pathways; and (C) Modularity (degree to which the network contains distinct subcommunities), which decreased dramatically in 2009, signifying breakdown of previously isolated virus-host relationship clusters. Solid black lines with points represent annual values calculated after aggregating edges by maximum weight between each virus-host pair. The red dashed vertical line indicates 2009, when the spiny water flea was officially detected.

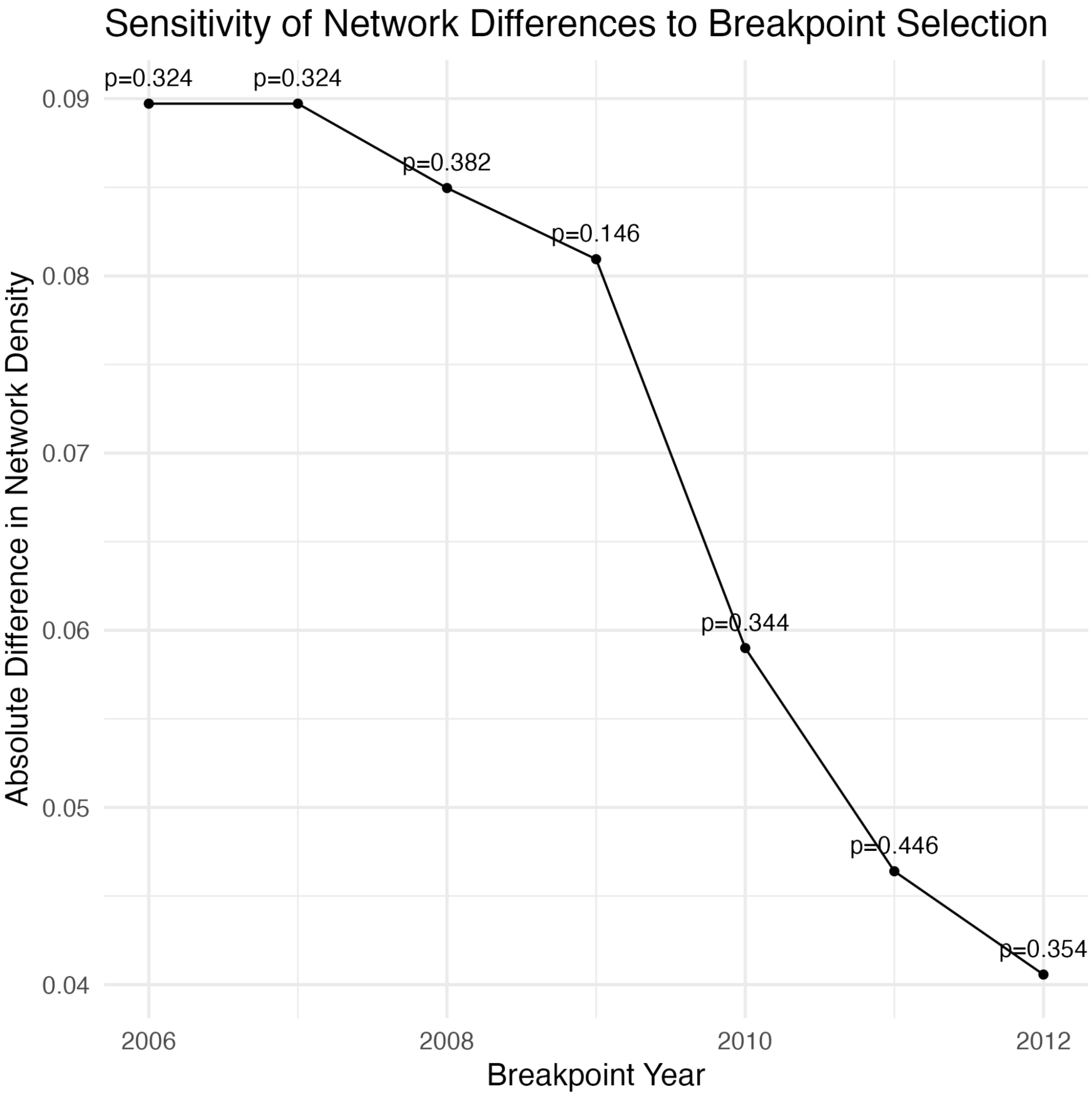

Supplementary Figure S9: Sensitivity analysis of network structural changes using multiple potential breakpoints for the spiny water flea invasion.
Each point represents a different analytical division of the dataset (2006-2012), with the x-axis indicating the tested breakpoint year and the y-axis showing the absolute difference in network density between pre- and post-breakpoint periods. P-values (displayed above each point) were calculated using permutation tests (500 permutations) that randomly reassigned samples to periods while maintaining original sample sizes.

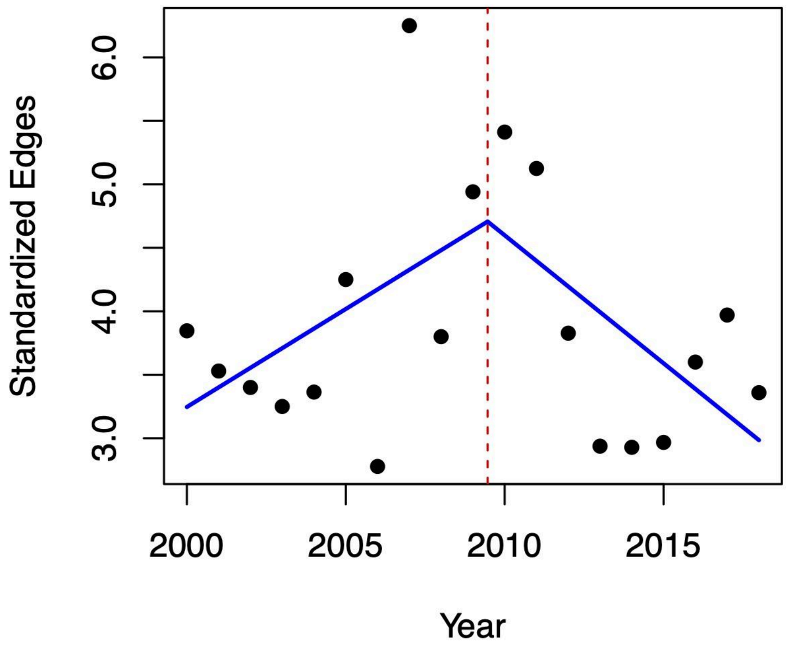

Supplementary Figure S10: Segmented regression analysis identifying a breakpoint in network edges over time.
Annual standardized network edges (edges per unit sampling effort) from 2000 to 2018 are shown as points. The solid blue lines represent segmented regression model fits before and after the estimated breakpoint, while the vertical dashed red line indicates the breakpoint estimate (year = 2009.5, 95% CI: 2004.6–2014.3). Although the breakpoint aligns closely with the documented invasion detection year (2009), slopes before and after this breakpoint were not statistically significant, suggesting that ecological changes associated with the invasion were gradual rather than abrupt.

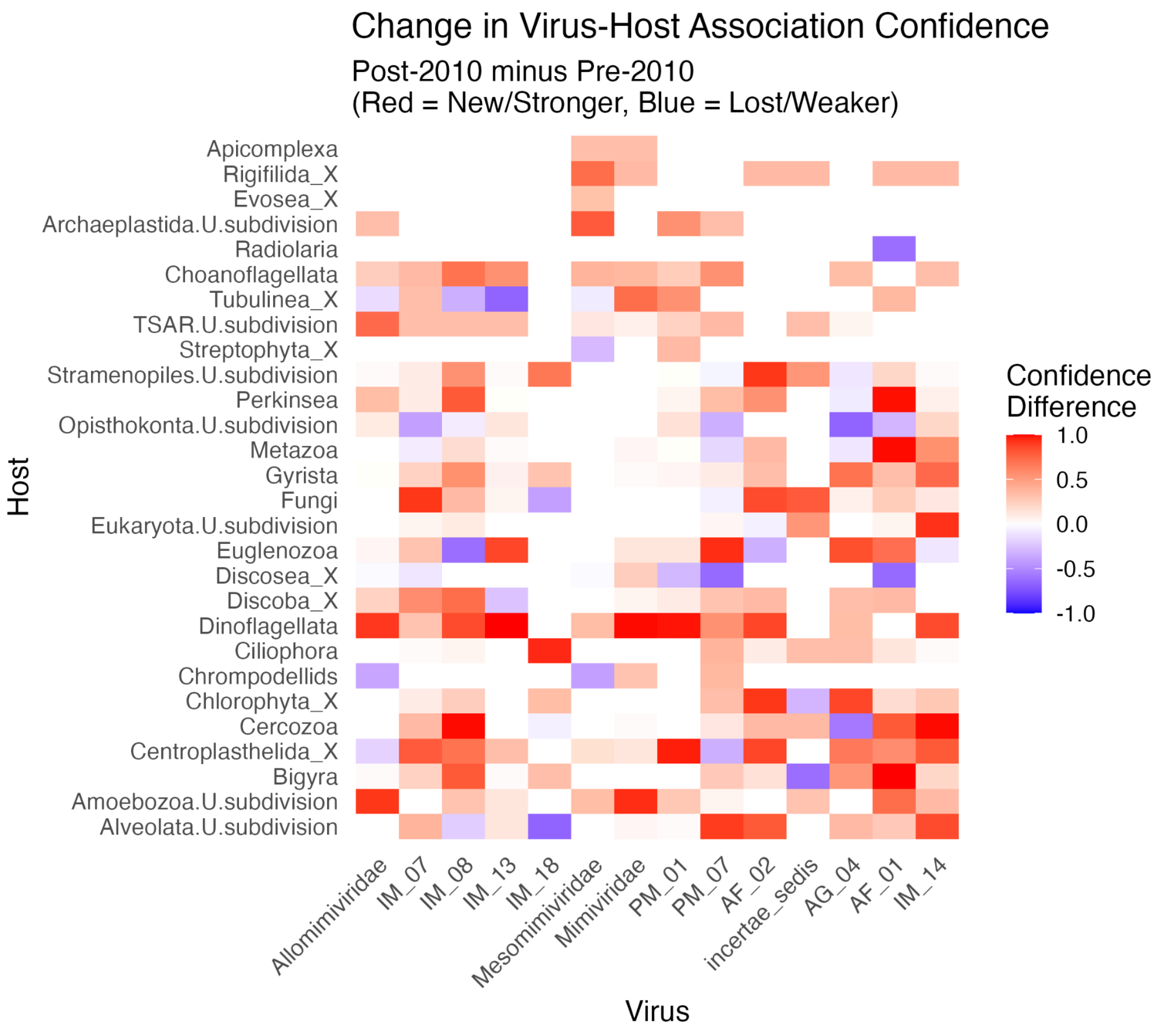

Supplementary Figure S11: Temporal shifts in virus-host association confidence between pre-2010 and post-2010 periods.

Heatmap displaying changes in virus-host association confidence between pre-2010 and post-2010 periods, where red indicates associations that strengthened or emerged after 2010 and blue shows associations that weakened or disappeared. Color intensity represents the magnitude of confidence change between periods (ranging from -1 to +1), with darker colors indicating stronger shifts. White regions indicate either consistently absent associations or those with unchanged confidence levels across both time periods.

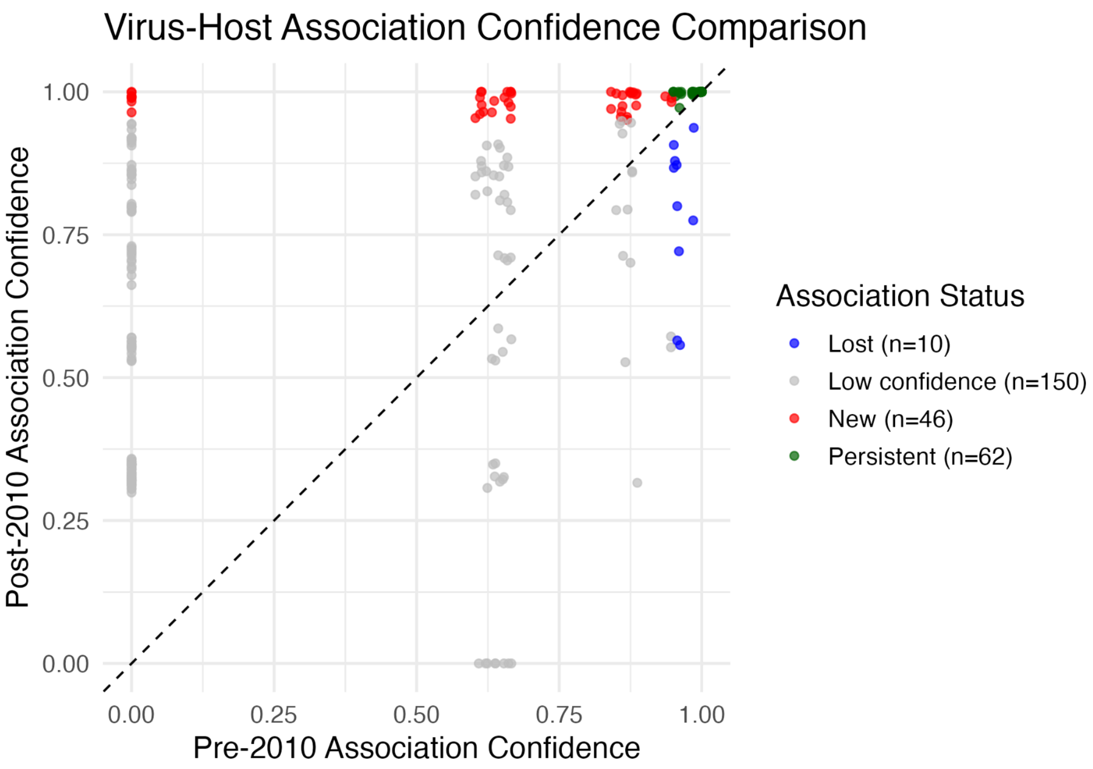

Supplementary Figure S12: Temporal comparison of virus-host association confidence scores between pre-2010 and post-2010 datasets.
Scatter plot comparing the bootstrap confidence values for all virus-host associations before (x-axis) and after (y-axis) 2010, with colors indicating association status: red points represent new high-confidence associations (≥95% in post-2010 only), blue points show lost associations (≥95% in pre-2010 only), green points indicate persistent associations (≥95% in both periods), and gray points represent low-confidence associations. Points above the dashed diagonal line represent strengthened associations following the spiny water flea invasion, while points below indicate weakened relationships.

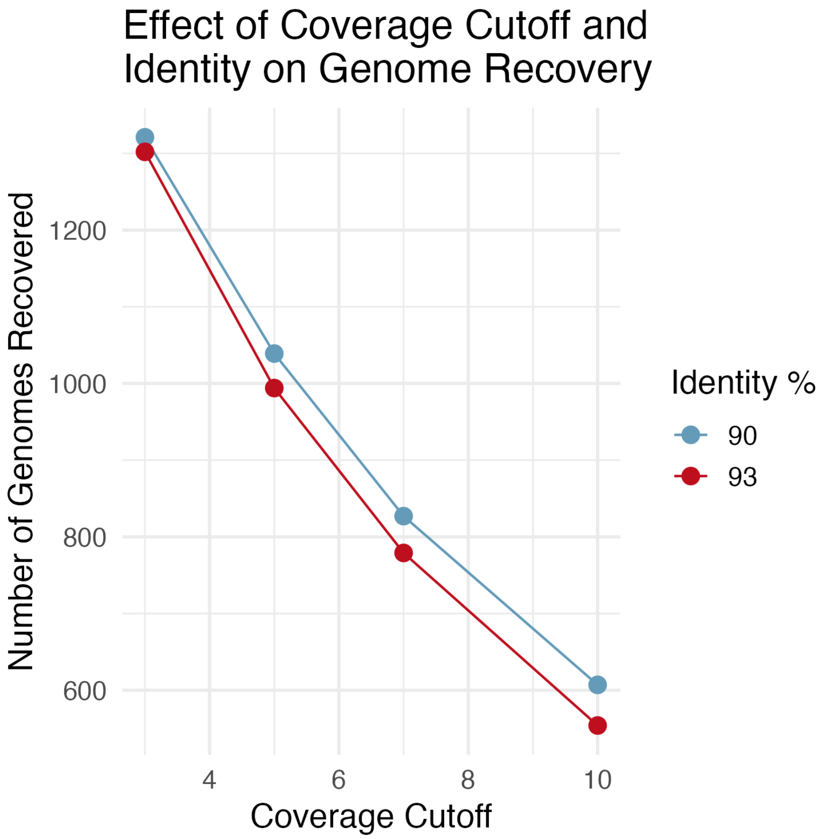

Supplementary Figure S13: Number of GVMAGs recruited by MetaPop with different percentages of minimum identity and coverage cutoff from MetaPop.
Coverage cutoff is highly significant ($p < 0.001$) with a coefficient of -102.95, with the lowest cutoff (cov = 3) yielding over twice as many genomes as the strictest cutoff (cov = 10). This indicates that for each decrease in coverage cutoff, we recovered ~103 more genomes. However, identity percentage is not statistically significant ($p = 0.35$). Thus, we keep our stricter percent identity and coverage cutoff to ensure false positives are not included.

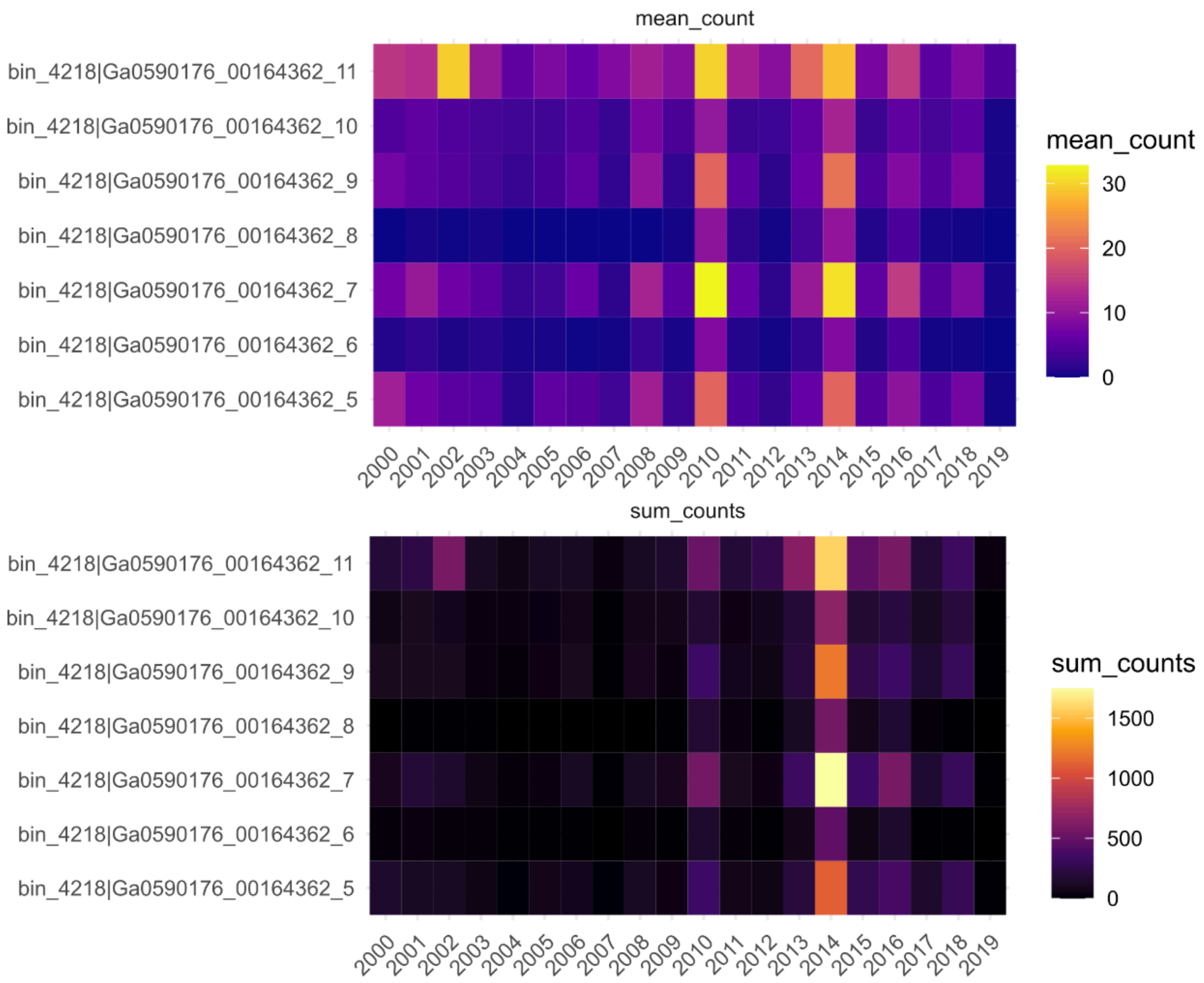

Supplementary Figure S14: Counts of phoH gene gain for all years.

It's important to note that our standard methods (shown in Figure 2B) could not detect gene 8 (our gene of interest) in samples from 2002-2008. To conclusively determine whether gene 8 was truly absent during these years, we implemented a more sensitive approach by combining all sequencing reads from each year and performing targeted read mapping specifically for the contig containing gene 8. From these results, we created mean count and sum count plots. This targeted approach provided higher sensitivity than our original method, clearly showing consistently low or absent counts of gene 8 during 2002-2008, followed by its sudden appearance around 2010. These findings confirm that the emergence of gene 8 represents a genuine gene gain event in the viral population rather than a sampling artifact or methodological limitation.
